# Human polymerase theta helicase positions DNA microhomologies for double-strand break repair

**DOI:** 10.1101/2024.04.26.591388

**Authors:** Christopher J. Zerio, Yonghong Bai, Brian A. Sosa-Alvarado, Timothy Guzi, Gabriel C. Lander

## Abstract

DNA double-strand breaks occur in all human cells on a daily basis and must be repaired with high fidelity to minimize genomic instability^1^. Deficiencies in high-fidelity DNA repair by homologous recombination lead to dependence on DNA polymerase theta, which identifies DNA microhomologies in 3’ single-stranded DNA overhangs and anneals them to initiate error-prone double-strand break repair. The resulting genomic instability is associated with numerous cancers, thereby making this polymerase an attractive therapeutic target^2,3^. However, despite the biomedical importance of polymerase theta, the molecular details of how it initiates DNA break repair remain unclear^4,5^. Here we present cryo-electron microscopy structures of the polymerase theta helicase domain bound to microhomology-containing DNA, revealing DNA-induced rearrangements of the helicase that enable DNA repair. Our structures show that DNA-bound helicase dimers facilitate a microhomology search that positions 3’ single-stranded DNA ends in proximity to align complementary base pairs and anneal DNA microhomology. We define the molecular determinants that enable the polymerase theta helicase domain to identify and pair DNA microhomologies to initiate mutagenic DNA repair, providing mechanistic insights into therapeutic targeting of these interactions.

## Introduction

Rapid and accurate repair of DNA double strand breaks is crucial for maintaining genome integrity. Since chromosome breaks, deletions, and rearrangements are enabling characteristics of cancer, robust and precise DNA repair mechanisms are necessary to minimize carcinogenesis^6^. The majority of DNA double-strand breaks are repaired with high fidelity through non-homologous end joining or homologous recombination (HR)^7,8^. Mutations in DNA damage repair genes cause increased dependency on an alternative, error-prone DNA repair pathway called theta-mediated end joining (TMEJ)^2,3^. These mutations are accompanied by an upregulation of DNA polymerase theta (Polθ), a multidomain protein that is critical for performing TMEJ^9,10^, and thereby associated with genome instability and carcinogenesis in HR-deficient cells^11-13^. Polθ is overexpressed in many cancers including breast, ovarian, lung, and colorectal cancers, and blocking Polθ expression or activity in the context of breast cancer-related gene (BRCA)1/2-deficient tumors is synthetic lethal^3,11,12^. While TMEJ allows cancer cells to repair DNA damage in an HR-deficient setting, this dependence on TMEJ also renders tumors particularly vulnerable to Polθ inhibition^2^. Thus, Polθ inhibitors hold therapeutic promise in combatting HR-deficient cancers^14-17^.

Polθ is a 290 kDa protein consisting of an N-terminal superfamily-2 helicase (PolθH) that is tethered to a C-terminal A-family polymerase (PolθP) via an unstructured 920-residue central domain (PolθC)^18^. Polθ engages and anneals DNA microhomologies two to six base pairs (bp) in length within resected 3’ single-stranded (ss)DNA overhangs at the site of a DNA double-strand break to initiate TMEJ^13,19,20^. Despite extensive study of this DNA repair system, the molecular mechanism by which Polθ identifies and anneals DNA micro-homologies is not well established^2,21^.

Both PolθP and PolθH are required for efficient TMEJ^22^, but the role of PolθH is poorly understood. Prior studies hypothesize that translocation of PolθH along ssDNA may remove DNA-binding proteins^23,24^, position PolθH to facilitate 3’ DNA microhomology annealing^21,23,25^, or suppress intrastrand base-pairing and snap-back replication by PolθP^22^. These mixed observations, along with the fact that PolθH displays weak DNA unwinding activity^26^, raise questions about the role of PolθH in 3’ DNA microhomology annealing and the interactions between PolθH and DNA that facilitate TMEJ. As a result, we sought to investigate the molecular determinants of the PolθH-DNA interaction.

## Results

### PolθH undergoes conformational rearrangements to facilitate microhomology annealing

To investigate the mechanistic relationship between PolθH and DNA microhomologies, we recombinantly expressed and purified human PolθH (residues 2-894, **Supplementary Fig. 1a**) for single particle cryo-electron microscopy (cryo-EM) analyses. In the absence of DNA, PolθH protomers assemble not only as a tetramer (**Fig. 1a**), which is consistent with prior crystallographic studies^25^, but also as a dimer (**Fig. 1b,c**), as recently observed for the cryo-EM structure of PolθH bound to inhibitor novobiocin^27^. Our *∼*3.3 Å resolution structure of the tetramer assembly adhered to D2 symmetry as a “dimer of dimers,” with an inter-dimer rotation of about 4 degrees compared to the tetramer crystal structure (**Supplementary Fig. 1b**)^25^. Our isolated dimer structure, resolved to *∼*3.5 Å resolution, is structurally consistent with both dimers that comprise the tetramer. Although neither of the previous PolθH structural studies detected a mixed population of assemblies^25,27^, we surmise that the dimeric PolθH may be the functional population, as the two helicase protomers are suited to handle the two resected ssDNA overhangs that are substrates for DNA repair by TMEJ.

**Figure 1.**
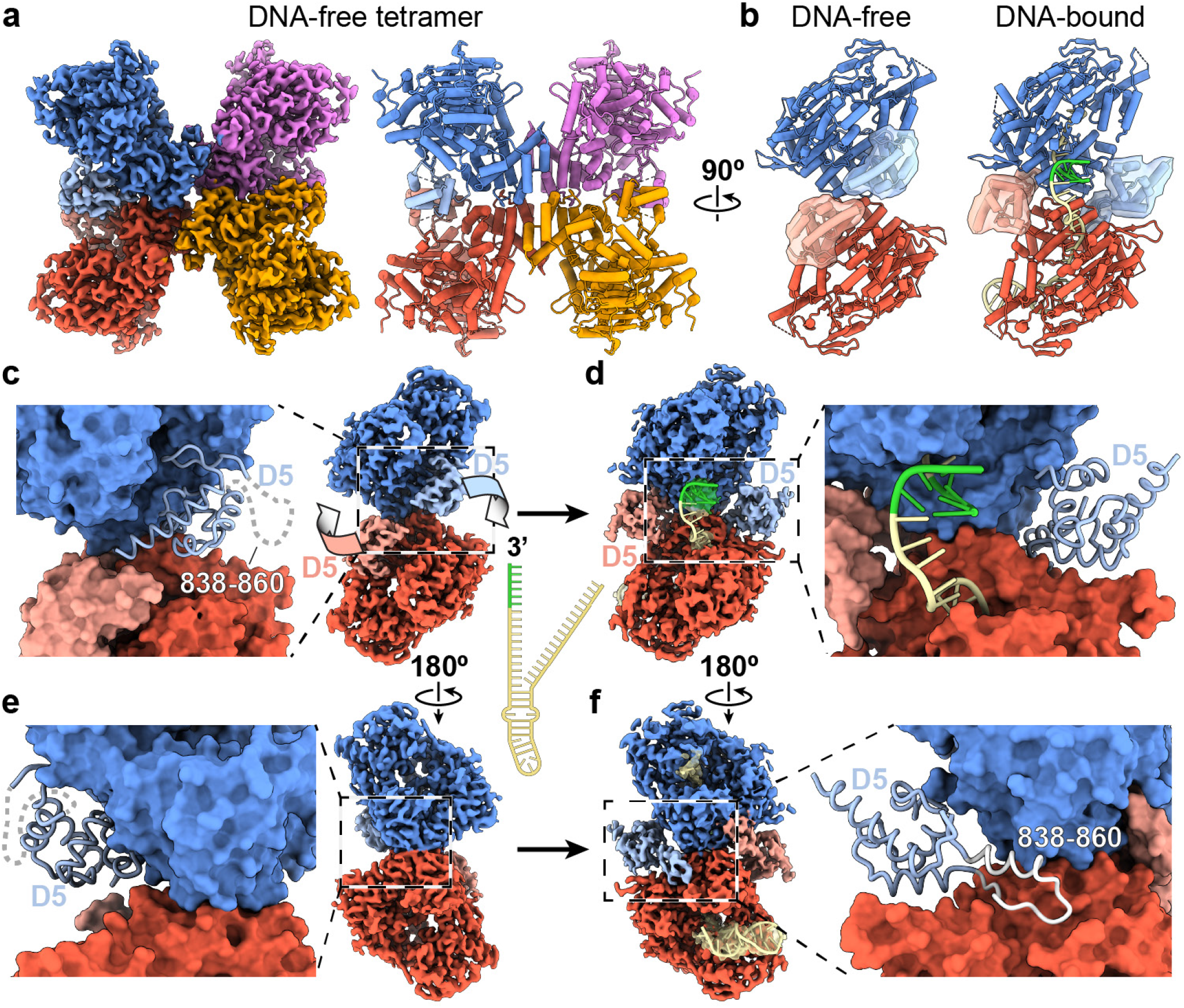
PolθH domain 5 rotates outward to accommodate DNA microhomology annealing. (**a**) Apo PolθH tetramer cryo-EM map and atomic model where helices are represented as tubes, colored by protomer. Domain 5 (D5) in each blue and red protomer is colored using a lighter shade. (**b**) Atomic models of the apo PolθH (left) and DNA-bound PolθH dimer (right) shown using a cartoon representation. The D5 in each protomer is highlighted using a semi-transparent surface. The annealed 3’ DNA microhomology (green) is shown between two helicase protomers in the DNA-bound dimer. (**c**) A close-up of the apo PolθH dimer atomic model is shown as a molecular surface, with D5 shown as a ribbon and the disordered residues 838-860 as a dashed gray line. On the right is the cryo-EM density of the apo PolθH colored as in (b), with arrows showing the trajectory of the D5 rearrangement that occurs upon interaction with DNA. A schematic of the DNA oligomer used for this study is shown to the right of the cryo-EM density, with the color green denoting the region of microhomology. (**d**) The cryo-EM density of PolθH bound to DNA is shown, with density corresponding to DNA extending from each DNA tunnel shown as a semi-transparent isosurface and the modeled DNA shown as a cartoon within (yellow with green microhomology). The closeup panel to the right emphasizes the new position of D5 after having rotated away from the center of the dimer to accommodate DNA pairing between protomers. (**e**) and (**f**) Views of the PolθH dimer maps, rotated 180*°* relative to the corresponding panels above in (c) and (d). The position of D5 before and after DNA binding is observable, with (f) emphasizing how repositioning of D5 causes residues 838-860 (white), which are disordered in the apo form, to adopt a loop-helix structure that docks into a pocket in the opposite protomer, stabilizing the DNA-bound dimer.

To gain further insight into PolθH function, we used a native PAGE assay to identify DNA species that bind PolθH. Our assay showed that PolθH preferentially binds DNA oligonucleotides with microhomology in a 3’ overhang (**Supplementary Fig. 2a,b**). This prompted us to introduce a stem-loop DNA with microhomology embedded in a 3’ over-hang to mimic the end-resected TMEJ DNA substrate. We expected this DNA substrate to interact with PolθH in a fashion that would recapitulate the initial stage of Polθ-mediated TMEJ with sufficient stability for cryo-EM structure determination.

We were immediately surprised to see in our cryo-EM images that the PolθH-DNA complex consisted entirely of dimers, indicating that binding of this DNA to PolθH drives dissociation of the tetrameric assembly. We determined a *∼* 3.5 Å resolution cryo-EM reconstruction of dimeric PolθH bound to DNA, enabling us to build an atomic model detailing the interactions between the helicase dimer and two strands of DNA that contain regions of 3’ microhomology (**Fig. 1b,d**). We identified density corresponding to ssDNA spanning the DNA tunnel of each protomer that was sufficiently resolved to discriminate between purines and pyrimidines. Following this ssDNA toward one of the 5’ ends, where one DNA strand presumably enters the tunnel, we observed density consistent with a double-stranded DNA stem loop. In the opposite direction, toward the center of the PolθH dimer, the 3’ end of the ssDNA exits the tunnel and connects to density corresponding to two 3’ ssDNA overhangs (one exiting from each PolθH protomer) annealed to form double-stranded DNA microhomology. This annealed microhomology is flexibly positioned above an open valley between the two PolθH protomers that lacks any stabilizing protein-DNA interactions, and thus could only be resolved at moderate resolution. However, given that the DNA density is connected to both incoming ssDNA oligonucleotides, and that the size of the structural feature can accommodate a span of 4-6 base-paired nucleotides, we are confident that the density between the protomers corresponds to the joined 3’ double-stranded DNA microhomology. No ATP was added to solve the PolθH-DNA structures, strongly indicating that PolθH ATPase activity is dispensable for PolθH identifying, aligning, and annealing 3’ DNA microhomology during TMEJ in the absence of ssDNA-binding proteins^21-23^.

We next compared our DNA-free and DNA-bound PolθH structures to investigate conformational changes associated with PolθH-DNA binding. This comparison revealed a striking rearrangement of the PolθH C-terminal domain (residues 790-894, hereafter referred to as D5). In our DNA-free PolθH structure, both D5s are positioned adjacent to each other in the intra-dimer valley, occupying the space where we observe DNA microhomology annealed in our DNA-bound structure (**Fig. 1c**). This adjacent positioning of the D5 domains at the protomer-protomer interface is consistent with the orientation of D5 in previous PolθH crystal structures^25^. However, in our DNA-bound structure, joining of DNA microhomology is associated with a repositioning of D5, whereby each D5 undergoes a *∼*150° rotation away from the dimeric axis, clearing the region between the two PolθH protomers to accommodate DNA microhomology annealing (**Fig. 1d, Supplementary Movie**). This D5-DNA relationship contrasts with the D5 equivalent in archaeal PolθH homolog *Archaeoglobus fulgidus* Hel308 (*∼* 30% sequence similarity to human PolθH), which directly interacts with ssDNA^28^. A more subtle and distinct D5 rearrangement was previously observed in the cryo-EM structure of PolθH bound to novobiocin^27^, suggesting that D5 is a mobile element that responds to or regulates functional states.

The large D5 rotation we observe is likely triggered by interactions formed by DNA as it traverses the PolθH DNA tunnel. We posit that as an incoming DNA strand exits the PolθH DNA tunnel, it encounters a negatively charged patch on D5 of the protomer, and resulting repulsive electrostatic interactions cause the outward swinging of D5 (**Supplementary Fig. 3**). This DNA-promoted D5 rearrangement establishes new intra- and inter-protomer interactions within the dimer, including the ordering of D5 residues 838-860, residues that were found to bind Rad51 recombinase in a previous study^12^, as a loop-helix that interacts with a pocket on the back side of the opposing protomer (**Fig. 1f, Supplementary Movie**). Conversely, in the DNA-free PolθH structure, these residues are flexible and not observed, and the interaction pocket on the back side of the dimer is not accessed (**Fig. 1e**). Formation of this loop-helix within this pocket on the opposite helicase protomer simultaneously locks D5 in the outward position to accommodate the annealed dsDNA while also stabilizing the inter-protomer interactions of the DNA-bound dimer.

The DNA-promoted rearrangement of D5 is concomitant with a larger scale rearrangement of the PolθH dimer that disrupts the tetrameric assembly (**Figs. 1f, 2a,b**). Upon DNA binding, the angle between the subunits of the PolθH dimer increases by *∼*14°, with the conserved protomer-protomer interface serving as a hinge point. Notably, the DNA-promoted rigid-body motions of the subunits, along with the new D5-mediated inter-protomer interactions, together destabilize the PolθH tetramer by sterically obstructing the inter-protomer interactions between the diagonal and adjacent protomers responsible for tetramer formation (**Fig. 2b**). These conformational changes of PolθH are consistent with a mechanism whereby PolθH binding DNA and DNA traversing the PolθH tunnel to the dimeric interface triggers a rearrangement of D5 that promotes tetramer dissociation, explaining why there are no tetrameric species in the DNA-bound PolθH population.

**Figure 2.**
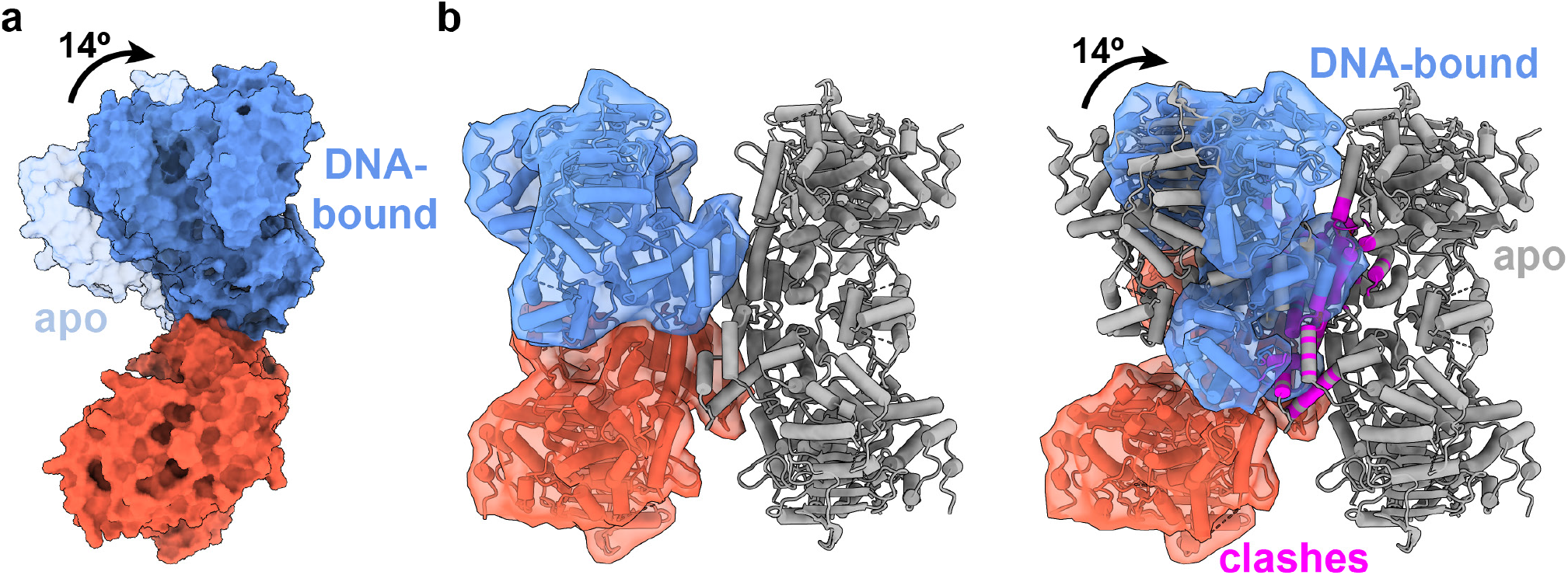
DNA binding to PolθH causes two protomers to hinge apart, destabilizing the PolθH tetramer. (**a**) Surface representations of the atomic models of the apo and DNA-bound PolθH dimers are shown after aligning the lower subunit (colored red) of each. The upper subunits are colored blue, with the apo subunit semi-transparent. Upon DNA-binding, the two PolθH protomers hinge 14*°* apart from each other. For clarity, DNA and D5 (residues 790-894) are omitted from each surface representation. (**b**) On the left, the apo tetramer is shown as a gray tube representation, with one dimer colored as in (a) and highlighted with a semi-transparent molecular surface. On the right, a DNA-bound dimer (colored and highlighted with DNA omitted) is overlaid on an apo tetramer (gray), aligned to the lower red protomer. The overlay demonstrates that upon DNA binding, the hinging and the D5 conformational change introduce steric clashes with the adjacent and diagonal tetramer protomers (clashing residues colored magenta), destabilizing the PolθH tetramer. Atoms less than 2 Å apart are considered to be clashing.

### DNA interactions with PolθH

Domains 1, 2, and 4 contribute to the DNA tunnel of PolθH (**Fig. 3a,b**), and the quality of our density in this region enabled us to identify key interactions between DNA and PolθH residues (**Fig. 3, Supplementary Fig. 4**). At the entrance of the DNA tunnel, three lysine residues (K348, K352, K497) are positioned for electrostatic interactions with the dsDNA backbone (**Fig. 3c**). Proximal to the tunnel entrance, a small loop in domain 2 comprising residues G465, G466, and R467 is positioned adjacent to the duplex DNA of the stem loop, at the branch point between the last paired DNA bases and the first unpaired bases of the 3’ DNA overhang (**Fig. 3d**). This nucleo-protein arrangement disrupts base pairing and stacking while directing the 3’ ssDNA overhang into the DNA tunnel, similarly to the corresponding β-hairpin DNA-unwinding loop in *A. fulgidus* Hel308 (**Supplementary Fig. 5a,b**)^28^. However, this loop is smaller in PolθH than in the Hel308 family helicases and lacks the aromatic residues that stack with the last of the paired DNA bases (**Supplementary Fig. 5b**)^28^. These differences may explain why PolθH does not possess the robust DNA unwinding activity displayed by *A. fulgidus* Hel308^2,26,29^.

**Figure 3.**
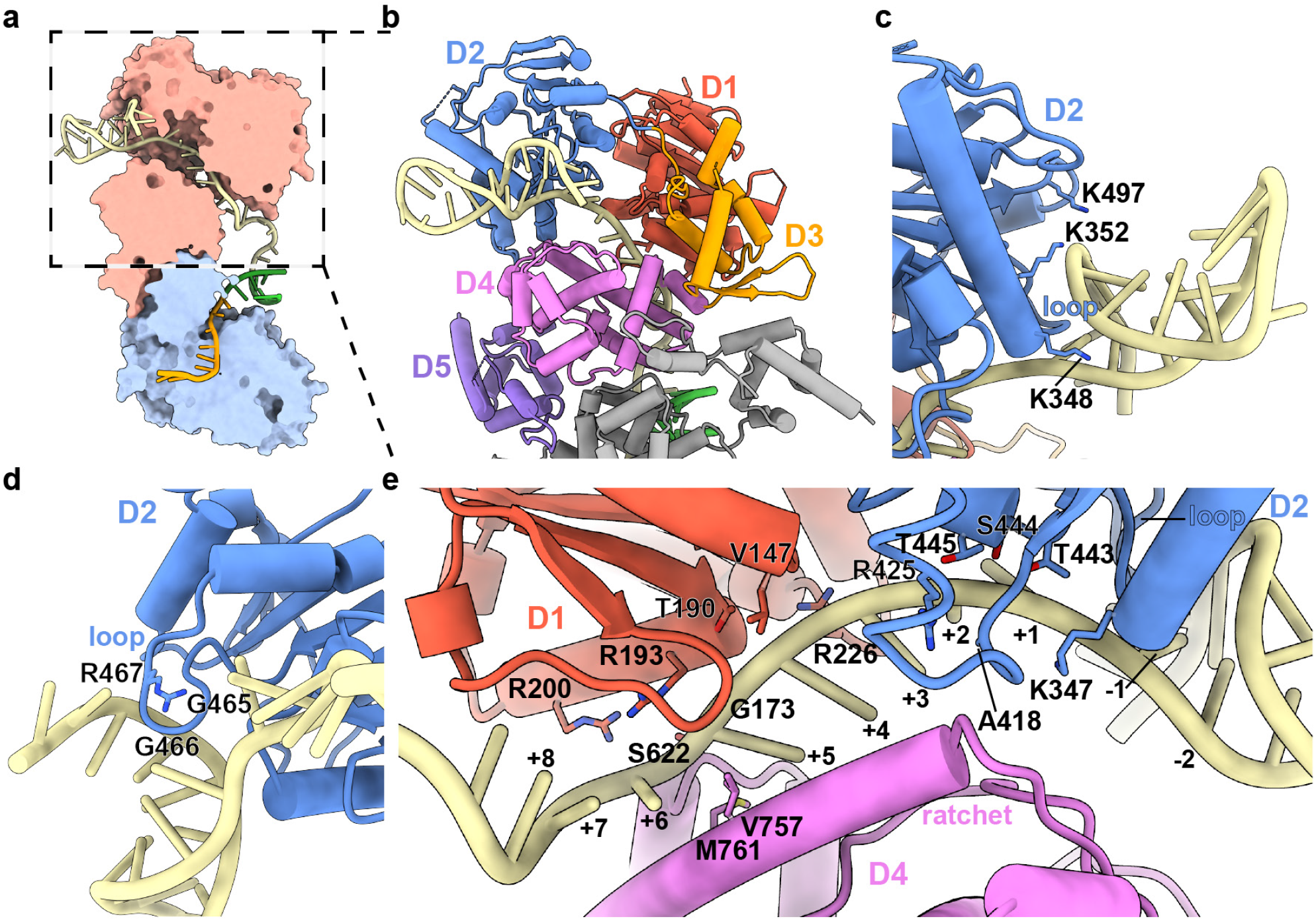
Characterization of PolθH-DNA binding. (**a**) Cross-section of the PolθH-DNA model showing that PolθH binds two copies of the DNA oligonucleotide (one orange and one yellow) and anneals the self-complementary 3’ microhomology (green). (**b**) Each domain of a single PolθH protomer is individually colored and labeled. (**c**) dsDNA interacts with residues in PolθH domain 2 adjacent to the ssDNA tunnel entrance. (**d**) A small PolθH loop facilitates ssDNA entry into the DNA tunnel. (**e**) DNA interacts with residues in PolθH domains 1, 2, and 4 as it traverses the DNA tunnel. DNA bases are labeled starting two bases upstream of the small PolθH unwinding loop from 5’ (−2) to 3’ (+8).

As ssDNA traverses the DNA tunnel, it contacts the ratchet helix - an ATP-coupled structural feature that has been shown to promote unidirectional (3’ to 5’) ssDNA translocation in Hel308 family helicases (**Fig. 3e**)^28,30^. In *A. fulgidus* Hel308, the ratchet helix ensures unidirectional ssDNA translocation via two residues (R592, W599) that stack with the bases of an incoming ssDNA substrate (**Supplementary Fig. 5c**)^28^. The PolθH ratchet helix, however, does not appear to introduce such base-stacking interactions with the DNA substrate, although residues V757 and M761 on this helix appear to wedge between two DNA bases, which may prevent back-tracking to facilitate processive 3’ to 5’ ssDNA translocation (**Supplementary Fig. 5d**)^24,25^.

The ssDNA is guided through the PolθH DNA tunnel via a series of base pair-agnostic interactions with the phosphate backbone (**Fig. 3e**). For clarity, we number the DNA bases from 5’ to 3’ with base -2 two bases upstream of the unwinding loop and base +8 exiting the DNA tunnel. Adjacent to the unwinding loop, K347 coordinates the phosphates of DNA bases +1 and +2, T443 and the backbone amine of A418 are positioned for hydrogen bonding interactions with the backbone of base +2, and T445 and R425 stabilize the backbone of DNA base +3. As the DNA continues through the tunnel, the DNA backbone approaches domain 1, where R226 and the backbone amine of V147 stabilize DNA base +4, T190 and the backbone amine of G173 stabilize +5, and R193 and R200 are positioned for ionic interactions with the backbone of base +6. As the DNA extends toward the tunnel exit, S622 stabilizes the base +6 backbone, and the bases point toward the domain 4 ratchet helix where V757 and M761 wedge between DNA bases +5 and +6 (**Supplementary Fig. 5d**). DNA bases +7 and +8 make no notable interactions with PolθH as they exit the DNA tunnel between domains 1 and 4.

### Multi-stage positioning of DNA overhangs for microhomology annealing

We can unambiguously trace ssDNA exiting the DNA tunnel from both PolθH protomers and joining as a large globular density in the intra-dimer valley. The location of this annealed DNA, combined with the external rotation of each D5 clearing a path for 3’ ssDNA overhangs to exit the DNA tunnels and extend toward one another, indicate that the intra-dimer valley serves as a nexus for microhomology pairing and associated allosteric rearrangements. Thus, we performed additional focused image analyses on this region, giving rise to PolθH reconstructions with 3’ DNA density in three distinct conformations between the PolθH protomers. The different positions of the DNA in these states present a putative three-step process of 3’ DNA microhomology pairing. In state 1, which we refer to as a microhomology “searching” state, we observe density corresponding to each 3’ overhang exiting the DNA tunnel and extending toward the first helix (residues 792-799) of the protomer’s D5 (**Fig. 4a**). In this configuration, the two ssDNA strands are positioned in an anti-parallel arrangement *∼*30 Å apart. This arrangement would enable ssDNA to simultaneously traverse each of the PolθH DNA tunnels and pass each other with the capacity to pair if there is complementary microhomology. Microhomology-driven interactions between the strands lead to state 2, which represents what we refer to as a “microhomology aligning” state (**Fig. 4b**). Here, both 3’ ssDNA overhangs are observed exiting the DNA tunnel and approaching D5, but then turn away from D5 toward each other to establish a nascent pairing in preparation for state 3, which we call the “microhomology annealed” state (**Fig. 4c**). In this state, the ssDNA moves away from D5 and the annealed microhomology forms a linear duplex between the DNA tunnel exits of both PolθH protomers, priming Polθ to continue TMEJ.

**Figure 4.**
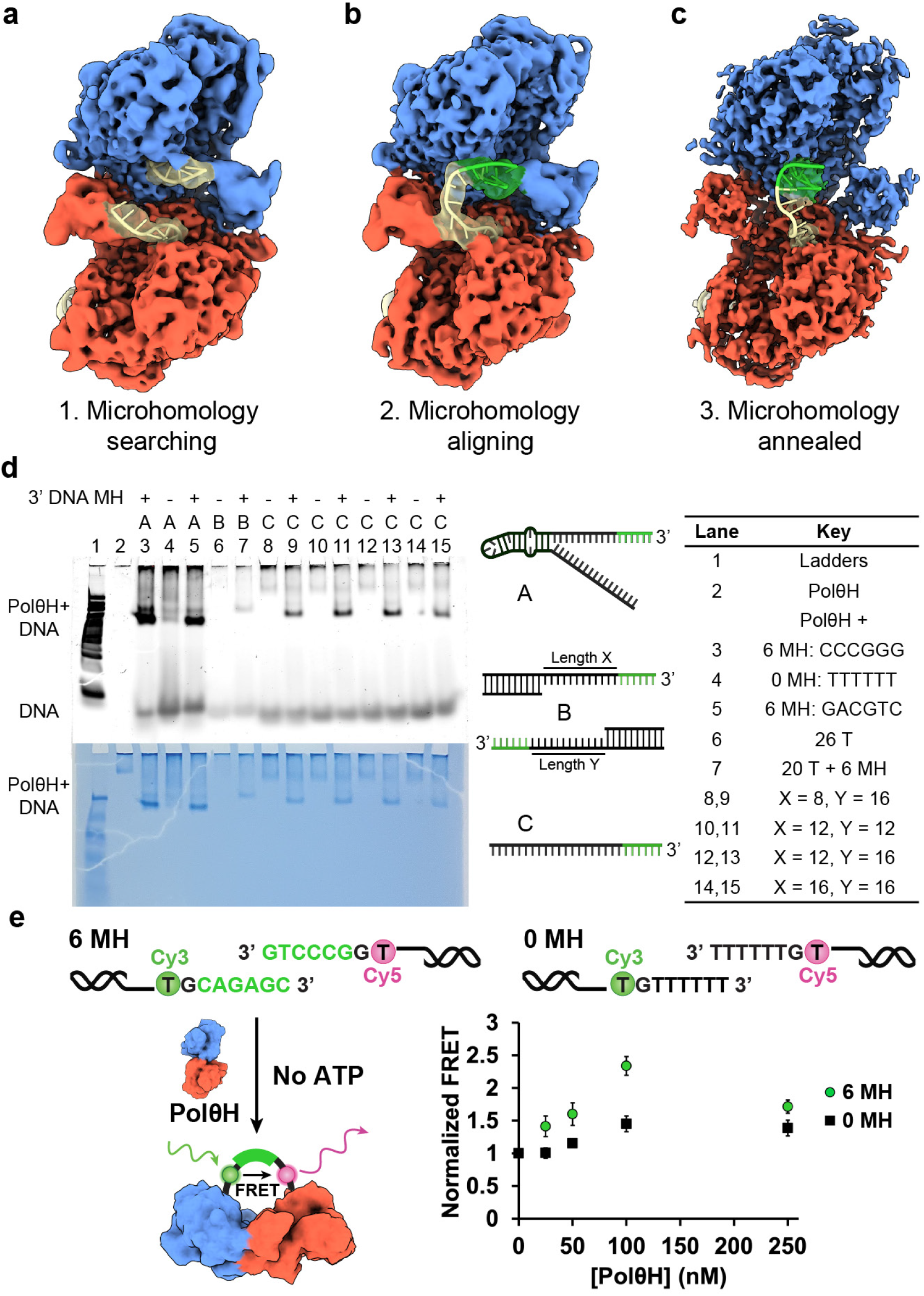
PolθH anneals 3’ DNA microhomology in 3 steps, as revealed by cryo-EM. Cryo-EM maps of DNA-bound PolθH shown in the 3’ microhomology (green) (**a**) searching, (**b**) aligning, and (**c**) annealed states. The cryo-EM density corresponding to the DNA extending from each DNA tunnel is shown as semi-transparent, with the modeled DNA shown as a cartoon within. The searching and aligning maps are shown locally filtered by resolution to emphasize DNA position. (**d**) Native PAGE assay illustrating that PolθH migrates more robustly into a native gel if there is 3’ DNA microhomology (MH) present. A key showing the types of oligonucleotides used is on the right. (**e**) FRET assay showing that PolθH brings 3’ DNA strands close together more effectively if there is 3’ DNA microhomology present. Error bars indicate standard deviation, n = 3.

Collectively, our structural data present a mechanism by which PolθH fosters ATP-independent 3’ ssDNA microhomology pairing and we sought to support this mechanism biochemically. We observed that PolθH alone, as well as PolθH incubated with DNA containing 3’ overhangs without microhomology, struggles to migrate into a native gel (**Fig. 4d**). Conversely, PolθH readily runs into a native gel upon addition of DNA containing 3’ overhangs with microhomology. We also noted that introducing 5’ secondary structure into the DNA results in a more stable interaction with PolθH, possibly because formation of a DNA stem loop prevents PolθH from fully translocating DNA, giving rise to a stalled complex. We concluded that PolθH can only form a stable complex and migrate into the native gel if there is 3’ microhomology present in the DNA substrate.

We next tested the capacity of PolθH to bring together 3’ ssDNA overhangs with (6 MH) and without (0 MH) microhomology by measuring FRET levels arising from PolθH-mediated joining of Cy3- and Cy5-labeled DNA substrates. PolθH facilitated increased FRET with both oligonucleotide pairs in a concentration-dependent manner in the absence of ATP, but we observed substantially higher FRET for the oligonucleotide pair containing 3’ microhomology (**Fig. 4e**). Notably, FRET increased with increasing concentrations of PolθH, but decreased at a protein concentration higher than 100 nM, indicating that at higher concentrations, PolθH dimers sequester individual free DNA ends, reducing the level of microhomology annealing^21^. Our observation that PolθH exhibits optimal ssDNA binding and annealing at concentrations ranging from 25-100 nM is consistent with previous fluorescence-based studies^21,22,25^. Our FRET data indicate that PolθH brings 3’ DNA ends together independent of DNA microhomology, but if the DNA overhangs contain microhomology, PolθH facilitates annealing, affording a more stable interaction consistent with a more robust FRET signal. Conversely, the minor FRET signal arising from the non-microhomology DNA likely corresponds to a population of PolθH in the DNA searching state, where the overhangs are brought in proximity but are unable to anneal. Together with our structural data, we conclude that the PolθH dimer functions as a platform capable of scanning ssDNA to identify,align, and anneal 3’ DNA microhomologies to initiate repair by TMEJ.

## Discussion

Cells carrying mutations in genes involved in DNA damage repair by HR, such as BRCA1/2, upregulate Polθ to repair DNA double-strand breaks via TMEJ, a process that is strongly correlated with HR-deficient cancers^12,31^. A crucial step in TMEJ is the Polθ-mediated pairing of 3’ ssDNA microhomologies. Our studies enable us to propose a mechanism by which PolθH searches 3’ DNA to identify, align, and anneal 3’ DNA microhomology without requiring ATP. Furthermore, our native PAGE and FRET assays show that the robustness of this activity is dependent on the presence of 3’ DNA microhomology.

In the context of prior studies, our findings present an updated mechanism for the initial steps of TMEJ (**Fig. 5, Supplementary Movie**). In the absence of DNA, PolθH exists in an equilibrium of tetrameric and dimeric states, both of which have accessible DNA tunnels for binding and initial translocation of the 3’ ends of resected ssDNA. Upon DNA binding and full traversal of the tunnel, interaction between the incoming ssDNA and the PolθH D5 C-terminal helix triggers a rearrangement of each D5 that stabilizes the PolθH dimer while simultaneously disrupting dimer-dimer interactions in the tetrameric form. These rearrangements are likely concomitant with further translocation of the ssDNA mediated by the interaction with D5, establishing the microhomology searching conformation observed in our cryo-EM data.

**Figure 5.**
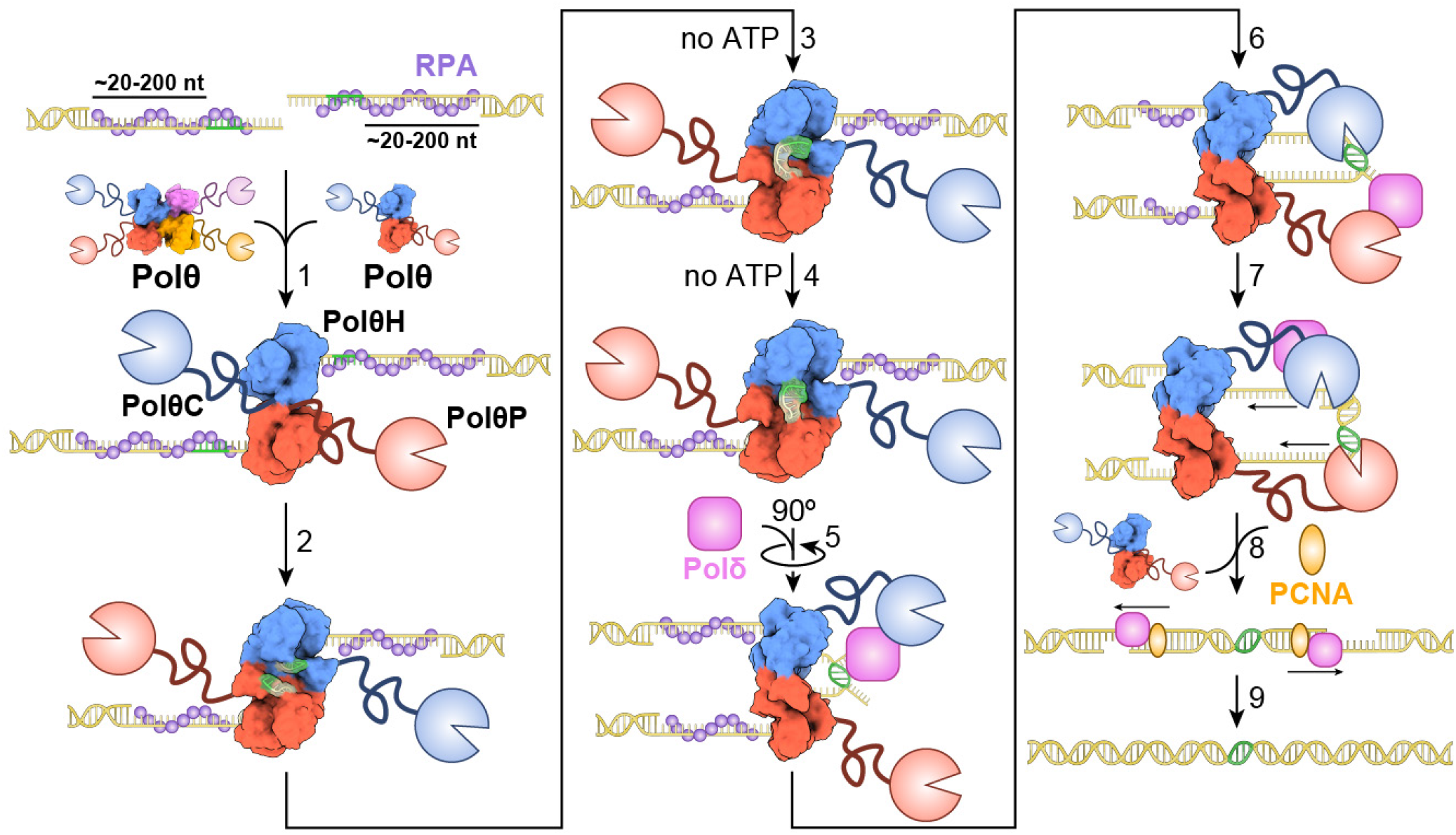
Proposed model for TMEJ, initiated by annealing of 3’ DNA microhomology by PolθH. Short-range DNA end-resection after a double-strand break leaves the TMEJ substrate with two 3’ ssDNA overhangs coated in ssDNA binding proteins, such as RPA. PolθH exists in equilibrium between tetrameric and dimeric forms before encountering 3’ ssDNA. (1) PolθH binds each 3’ ssDNA terminus. (2) PolθH translocates 3’ - 5’ on the DNA and removes RPA in an ATP-dependent manner. This DNA translocation facilitates PolθH domain 5 swinging open to accommodate the DNA. PolθH aligns (3) and anneals (4) 3’ DNA microhomologies in an ATP-independent manner. (5) Polδ trims one 3’ DNA flap and may stabilize the joined microhomology. (6) PolθP attaches to the available 3’ primer terminus formed by the joined microhomology and stabilizes it, and Polδ may trim the opposite 3’ DNA flap. (7) PolθP begins non-processive DNA synthesis. This may occur in only one direction, or the other copy of PolθP may begin non-processive DNA synthesis of the other side of the microhomology. (8) After minimal synthesis by PolθP, Polδ and PCNA perform processive, strand-displacement synthesis before (9) final resolution of the DSB.

We posit that this dimeric conformation enables 3’ to 5’ DNA translocation and displacement of replication protein A (RPA) from the ssDNA by PolθH in an ATP-dependent manner^23,24^. PolθH serves as a platform to bring DNA ends together, and given the 93% likelihood that a 3 bp microhomology is present within 15 bp of any given pair of DNA ends, DNA microhomology is likely identified and annealed almost immediately after DNA translocation begins^5^. However, given the limited number of base-pairing interactions, microhomology annealing is transient and prone to strand dissociation without stabilization by other TMEJ factors such as PolθP or DNA polymerase delta (Polδ)^21,32^. Thus, PolθH plays a key role in identifying and positioning regions of 3’ microhomology for subsequent annealing and stabilization for TMEJ, a conclusion that is supported by recent single-molecule FRET studies that characterized the contributions of PolθH and PolθP to TMEJ^21^. Mounting evidence suggests that after microhomology annealing, Polδ trims one unpaired 3’ ssDNA end (flap) and may bind Polθ in a manner that stabilizes the annealed microhomology^32^. After flap-trimming, one of the two tethered PolθPs may recognize a 3’ dsDNA microhomology primer, further stabilizing the annealed microhomology, and subsequently begin non-processive synthesis^21,33^. Once the first polymerase has gap-filled a sufficient distance from the microhomology, Polδ may trim the other unpaired 3’ ssDNA flap so that the second tethered copy of PolθP can begin synthesis on the opposite 3’ end of the microhomology. After less than 15 nt of synthesis by PolθP, Polδ can perform processive strand-displacement synthesis in complex with the proliferating cell nuclear antigen (PCNA) processivity clamp, triggering final resolution of the DNA break^32^.

In the context of this proposed mechanism, Polδ likely works in complex with PolθH and PolθP to stabilize the annealed microhomology, trim unpaired 3’ DNA flaps, and remain in proximity to perform DNA synthesis after PolθP^32^. Interaction with Polδ may also facilitate Polθ exit from the DNA substrate, likely before PolθH encounters 5’ dsDNA. This may explain why Polθ, with its small unwinding loop, has no need to act as a processive helicase. Our proposed mechanism also implicates a role for the disordered PolθC, which likely functions as a tether of defined length that maintains PolθP in proximity of PolθH to recognize paired microhomology while granting the polymerase the degrees of freedom necessary to robustly grasp, stabilize, and start non-processive synthesis. We anticipate that these insights into the molecular determinants of the PolθH-DNA interaction, which outline the critical steps required to initiate TMEJ, provide the mechanistic details to enable rational design of future PolθH-targeted cancer therapeutics.

## Materials and Methods

### Protein Expression and Purification

10xHis-SUMO-Avi-tagged human PolθH (amino acids 2-894) was expressed in *E. coli* Rosetta (DE3) cells in terrific broth. The culture was grown to an OD_600_ of 2.0 at 37 °C, transferred to 16 °C, and induced with 0.5 mM IPTG for 20 hrs. Cells were pelleted and resuspended in lysis buffer (40 mM Hepes pH 7.5, 0.5 M NaCl, 5% glycerol, 0.5 mM TCEP, 1 mM PMSF), and lysed by sonication. The lysate was clarified by centrifugation, and the supernatant was incubated with Ni-NTA resin (Qiagen) at 4 °C for 2 hrs. The mixture was applied to a gravity column, the beads were washed with lysis buffer and lysis buffer plus 20 mM imidazole, and the protein was eluted with lysis buffer plus 250 mM imidazole. The eluted protein was digested with His-tagged TEV protease during dialysis in lysis buffer at 4 °C for 12 hrs to remove the His-SUMO-Avi tag. The protein mixture was incubated on Ni-NTA resin and loaded onto a gravity column, the flowthrough was collected, and the resin was washed with dialysis buffer.

PolθH was concentrated and loaded onto a 5 mL HiTrap Heparin column (Cytiva). The column was washed with Heparin buffer A (50 mM Hepes pH 7.5, 100mM NaCl, 5% glycerol, 0.5 mM TCEP), PolθH was eluted from the column with a gradient of 0 - 50% Heparin buffer B (50 mM Hepes pH 7.5, 1M NaCl, 5% glycerol, 0.5 mM TCEP), and the column was washed with 100 % Heparin buffer B. Fractions containing PolθH were pooled, concentrated, and further purified by size exclusion chromatography using a Superdex 200 10/300 GL (Cytiva) column and SEC buffer (10 mM Hepes pH 7.5, 250 mM NaCl, 0.5 mM TCEP). PolθH was collected and subject to endotoxin removal with a ToxinEraser Endotoxin Removal kit (GenScript). PolθH was concentrated, aliquoted, flash-frozen, and stored in SEC buffer at -70 °C.

### Native PAGE

All DNA oligonucleotides used in this study are shown in **Supplementary Table 1**. Frozen PolθH protein in storage buffer (10 mM Hepes-NaOH, pH 7.5, 250 mM NaCl, 0.5 mM TCEP) was thawed on ice, and 2*×* pre-annealed oligonucleotides (IDT) were added. The mixture was incubated on ice for 20 min, NaCl concentration was lowered to 100 mM over the course of 1 hr with addition of no-salt storage buffer (10 mM Hepes-NaOH, pH 7.5, 0.5 mM TCEP), and the mixture incubated on ice for 20 min. Glycerol was added to a final concentration of 5% and samples were loaded on a 4.5% Native PAGE gel that had pre-run at 75 V for 90 min in 0.2× TBE buffer pH 7.6 at 4 °C. The gel ran at 4 °C for 90 min, was rinsed with water, and then placed in a 50 mL solution of 1 *×*Diamond Nucleic Acid Dye (Promega) in 0.2 *×*TBE buffer pH 7.6 to rock in the dark for 30 min. DNA bands were visualized on a ChemiDoc MP imaging system (Bio-Rad). The gel was then rinsed with water, incubated in Coomassie protein stain, de-stained, and protein bands were visualized on a lightbox.

### Preparation of PolθH-DNA complex

PolθH was thawed on ice, and pre-annealed 2*×* oligonucleotides (IDT) were added. The mixture incubated on ice for 30 min, NaCl concentration was lowered to 100 mM over the course of 1 hr with addition of no-salt storage buffer (10 mM Hepes-NaOH, pH 7.5, 0.5 mM TCEP), and the mixture incubated on ice for 30 min. The resulting complex was loaded onto a Superdex 200 10/300 GL (Cytiva) column pre-equilibrated with low-salt SEC buffer (10 mM Hepes-NaOH, pH 7.5, 100 mM NaCl, 0.5 mM TCEP), for size-exclusion chromatography at a flow rate of 0.5 ml/min. Fractions containing protein were collected and analyzed using native PAGE. Fractions with both protein and DNA were collected and concentrated. Concentration was determined using a BCA assay and samples were immediately prepared for cryo-EM.

### Preparation of samples for cryo-EM

Samples were kept at 4 °C throughout the sample preparation process. Apo PolθH was diluted to 0.9 mg/mL in SEC buffer, PolQHel-DNA complex was diluted to 1.25 mg/mL in low-salt SEC buffer, and samples were subject to centrifugation at 15,000 *×* g for 10 min. Quantifoil 300 mesh R 1.2/1.3 Holey Carbon Films were glow discharged under vacuum for 30 s at 15 mA in a Pelco easiGlow 91000 Glow Discharge Cleaning System (Ted Pella, Inc.). The sample (3 μL) was applied to the surface of the grid, blotted with Whatman 1 filter paper until 2 s after the liquid spot on the filter paper stopped spreading, and the grid was then immediately plunged into a liquid ethane pool cooled by liquid nitrogen using a manual plunge freezer in a 4 °C cold room with *>*95% humidity. Grids were then clipped under liquid nitrogen in preparation for imaging.

### Cryo-EM data acquisition

Cryo-EM data were acquired using a Titan Krios transmission electron microscope with a field emission gun operating at 300 keV. Movies were collected using a Gatan K3 BioQuantum direct electron detector with the energy filter set to 20 eV, operated in electron counting mode. The Leginon data collection software (v3.6)^34^ was used to collect micrographs at 105,000 *×* nominal magnification (0.833 Å /pixel at the specimen level) with a nominal defocus set to 1.0 μm under focus. Apo PolθH movies were collected with the slit out, at an exposure rate of 38.8 e^-^/pixel/s, with 30 frames (30 ms each) over 0.9 s, for a total electron exposure of 50 e^-^/Å ^2^. PolθH-DNA movies were collected with the slit in, at an exposure rate of 24.8 e^-^/pixel/s, with 47 frames (30 ms each) over 1.4 s, for a total electron exposure of 50 e^-^/Å ^2^. Stage movement was used to target the center of four 1.2 μm holes for focusing and an image shift was used to acquire high magnification images in the center of a larger array of targeted holes. Detailed dataset-specifics are available in **Supplementary Table 2**.

### Cryo-EM data analysis of apo PolθH

The image processing pipeline for the apo dataset is shown in **Supplementary Figure 6**. Frames from cryo-EM movies were aligned and combined applying a dose-weighting scheme using MotionCor2^35^ using default settings and transferred to cryoSPARC live v4.2.1 - 4.3.0^36^ for CTF estimation with patchCTF using default parameters. 4,164 micrographs were collected, micrographs with reported CTF resolutions lower than 6.5 Å were discarded, and 3,906 micrographs were exported to cryoSPARC v4.2.1-4.3.0.^37^ Particles were picked using a blob picker, and an initial round of 2D classification was used to generate templates for template picking with a 130 particle diameter and 0.7 *×* diameter minimum separation distance. 1,643,557 particles were picked using template-based picking and 1,474,207 particles were extracted with a 320-pixel box without any Fourier binning. All particles were subject to minimal cleanup based on 2D classification using 70 classes, and 4 initial models were generated Ab initio from 1,436,257 particles contributing to selected class averages showing detailed secondary structure.

Tetrameric, dimeric, and junk populations were identified from the same population of selected particles using two rounds of heterogeneous refinement (401,032 tetramer and 399,289 dimer particles). Each population was refined using non-uniform refinement using default parameters with and without symmetry applied (tetramer D2, dimer C2).^38^ Symmetry was validated by rigid body fitting the corresponding C1 maps into the symmetrized maps in ChimeraX (UCSF)^39,40^. The symmetry-imposed dimer and tetramer particles were symmetry expanded (1,604,128 tetramer protomer particles and 798,578 dimer protomer particles) and a mask around a single protomer in each structure was used to perform a local refinement with rotation and shift search extents of 1 degree and a maximum alignment resolution of 0.1 degree to improve the resolution of the protomer. The refined tetramer protomer density was further refined using 3D variability analysis in cluster mode using 10 clusters to remove junk protomers. 1,233,292 intact tetramer protomers were subject to 3D classification without alignment using 3 classes, 8 Å target resolution, O-EM learning rate init 0.2, PCA initialization, forced hard classification, and class similarity 0.1, and then 3D variability analysis in cluster mode using 10 clusters.

The refined dimer protomer density was further refined using 3D variability analysis in cluster mode with 5 clusters. A small subset of protomer particles from each population was chosen to generate final protomer maps through masked local refinement based on homogeneity across the entire particle, quality of map features, and estimated resolution by FSC at a cutoff of 0.143. For each population, a sharpened protomer map and a protomer mask was opened in ChimeraX and command “volume multiply” was performed. The resulting protomer maps were copied and fit into corresponding symmetric maps in ChimeraX, and command “volume resample” on the combined map grid was used to produce a combined tetramer map and a combined dimer map. All map-based parameters for the apo dataset refer to the locally refined individual protomer maps. Sphericity and resolution of protomer maps were evaluated by the remote 3DFSC processing server and PHENIX mtriage^41,42^. Local resolution plots were generated within cryoSPARC for each reconstruction and are found in **Supplementary Figure 8**. These plots were used to determine local resolution ranges of both protomer maps using “Report value at mouse position” within the ChimeraX “surface color” tool.

### Cryo-EM data analysis of apo PolθH-DNA

The PolθH-DNA dataset image processing pipeline is shown in **Supplementary Figure 7**. Frames from cryo-EM movies were aligned and combined applying a dose-weighting scheme using MotionCor2^35^ and transferred to cryoSPARC Live v4.2.1 - 4.3.0^36^. 12,440 micrographs were collected, CTF estimation was performed with live patchCTF using default parameters, and micrographs with reported CTF resolutions lower than 5.7 Å were discarded, leaving 12,201 micrographs. Particles were picked using a blob picker, and an initial round of live 2D classification was used to generate templates for template picking with a 120 Å particle diameter and 0.7 *×* diameter minimum separation distance. 5,424,691 particles were picked using template picker, extracted with a 324-pixel box without any Fourier binning, and exported to cryoSPARC for minimal cleanup based on 2D classification with 120 classes. Three maps were generated Ab initio from a subset of the 5,050,169 particles contributing to selected class averages, and one was chosen as an initial model based on presence of DNA density between two PolθH protomers. This initial model, along with trash classes, were used to seed iterative heterogeneous refinement of all quality particles to remove classes with no density for DNA or domain 5. The remaining 1,111,072 particles were subject to 3D variability analysis in cluster mode using 10 clusters, which revealed different DNA positions among PolθH-DNA classes. 413,973 particles in the microhomology annealed state were subject to an additional round of 3D variability analysis in cluster mode using 8 clusters and then masked 3D classification without alignment using 2 classes, 10 Å target resolution, O-EM learning rate init 0.2, PCA initialization, forced hard classification, and class similarity 0.1 to identify a population with the strongest density for annealed microhomology.

Populations in non-annealed DNA states were combined and subject to 3D variability analysis in cluster mode using 5 clusters and then heterogeneous refinement to separate populations in DNA searching and aligning states. Subsets of particles in each microhomology pairing state were chosen as final models selected for homogeneity across the entire particle, quality of map features, and estimated resolution by FSC at a cutoff of 0.143. Per-particle defocus and up to fourth-order optical aberrations were corrected in cryoSPARC’s non-uniform refinement protocol to generate sharpened maps of each particle population. Local resolution plots were generated within cryoSPARC for each reconstruction and are found in **Supplementary Figure 8**. These plots seeded cryoSPARC local filtering jobs to produce maps that were locally filtered by resolution and were used to determine local resolution ranges of each sharpened map using “Report value at mouse position” within the ChimeraX Surface Color tool. Sphericity and resolution of protomer maps were evaluated by the Remote 3DFSC Processing Server and PHENIX mtriage^41,42^.

### Cryo-EM data analysis of apo PolθH-DNA

Model building and refinement were initiated with published models for PolθH (PDB: 5A9J)^25^ and *A. fulgidus* Hel308 (PDB: 2P6R)^28^. For the apo reconstructions, one protomer was modeled into one protomer map and the model was symmetry expanded to fit each combined map in ChimeraX. The combined models were then further refined in their respective combined maps. Iterative rounds of model building and refinement were performed in Coot v0.9.8.1 EL^43^, ISOLDE v1.6.0^44^, and PHENIX v1.20.1^45^ until reasonable agreement between the model and data were achieved. Finally, after Ramachandran parameters, rotamers, and clashes were satisfied in ISOLDE, ISOLDE command “isolde write phenixRsrInput #*<*model*> <*map resolution*>* #*<*map*>*” was used to export the model and generate a rigid-body refinement settings file for PHENIX real-space refine. PHENIX real-space refinement was performed with the output model, rigid-body settings file, and sharpened map, and PHENIX Comprehensive validation (cryo-EM) was performed using the map and output model to yield map and model parameters. Quality of model fitting to map is evaluated in per-residue correlation coefficient (CC) outputs from Phenix as shown in **Supplementary Figure 9**, as well as scores from MolProbity^46^, EMRinger^47^, and Q-score^48^ shown in **Supplementary Tables 2**. Q-scores were calculated with σ = 0.4. ChimeraX was used to interpret the EM reconstructions and atomic models, as well as to generate figures.

### FRET

Frozen PolQHel protein in storage buffer (10 mM Hepes-NaOH, pH 7.5, 250 mM NaCl, 0.5 mM TCEP) was thawed on ice. Individually annealed Cy3 and Cy5 oligonucleotides (IDT) were mixed in a 1:1 ratio, protein was added, and the samples incubated on ice for 20 mins. FRET buffer (25 mM Hepes NaOH pH 7.5, 100 mM NaCl, 1mM MgCl2, 0.01% NP-40, 5% glycerol, 2 mM DTT) was added to dilute the sample to the indicated protein concentration and 25 nM FRET oligonucleotide substrate, and the samples equilibrated at room temperature for 20 mins. 50 μL of each sample was added to a black 384-well plate (Greiner Bio-One 781900). FRET (excitation: 550 nM, emission: 668 nM) was read on a Biotek Synergy H1 microplate reader (Agilent) with Gen5 software v3.02.

## Acknowledgements

We thank JC Ducom at Scripps Research High Performance Computing and Charles Bowman at Scripps Research for computational support, as well as Will Lessin at the Scripps Research Electron Microscopy Facility for microscopy support. Research reported in this publication was supported by the National Cancer Institute of the National Institutes of Health (NIH) under Award Number F32CA288144 (CJZ), GCL is supported by NIH grant GM14305, and the work used equipment supported by NIH grant S10OD032467.

## Data availability

Cryo-EM maps and associated atomic models were deposited to the Electron Microscopy Databank (EMDB) and the Protein Databank (PDB), respectively, with the following EMDB and PDB IDs: apo PolθH tetramer - EMD-44534, 9BH6; apo PolθH dimer - EMD-44535, 9BH7; PolθH-DNA microhomology searching - EMD-44536, 9BH8; PolθH-DNA microhomology aligning - EMD-44537, 9BH9; PolθH-DNA microhomology annealed - EMD-44538, 9BHA.

## Author Contributions

C.J.Z. prepared all cryo-EM samples, collected data, produced high-resolution structures, built all atomic models, performed all biochemical experiments, and wrote the manuscript. Y.B., B.A.S., and T.G. provided initial support on PolθH-DNA binding studies and the native PAGE assay. G.C.L. and C.J.Z. designed all experiments and performed all mechanistic interpretation. G.C.L. provided guidance in cryo-EM data collection, analyses, and edited the manuscript.

## Supplementary Figures

**Supplementary Figure 1.**
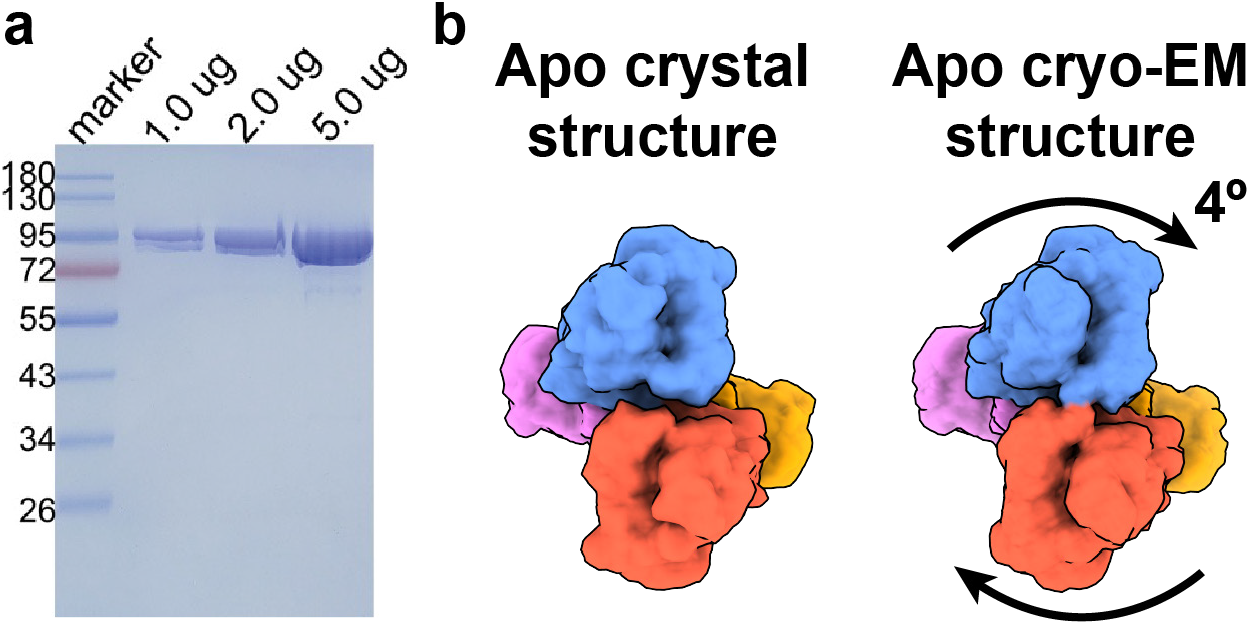
Comparison of apo PolθH tetramer cryo-EM and crystal structures. (**a**) Coomassie-stained SDS-PAGE of PolθH residues 2-894. (**b**) In the PolθH tetramer cryo-EM structure, two dimeric PolθH subunits are rotated about 4 degrees with respect to the crystal structure (PDB: 5A9J). Residues 35-66 are omitted from the crystal structure representation.

**Supplementary Figure 2.**
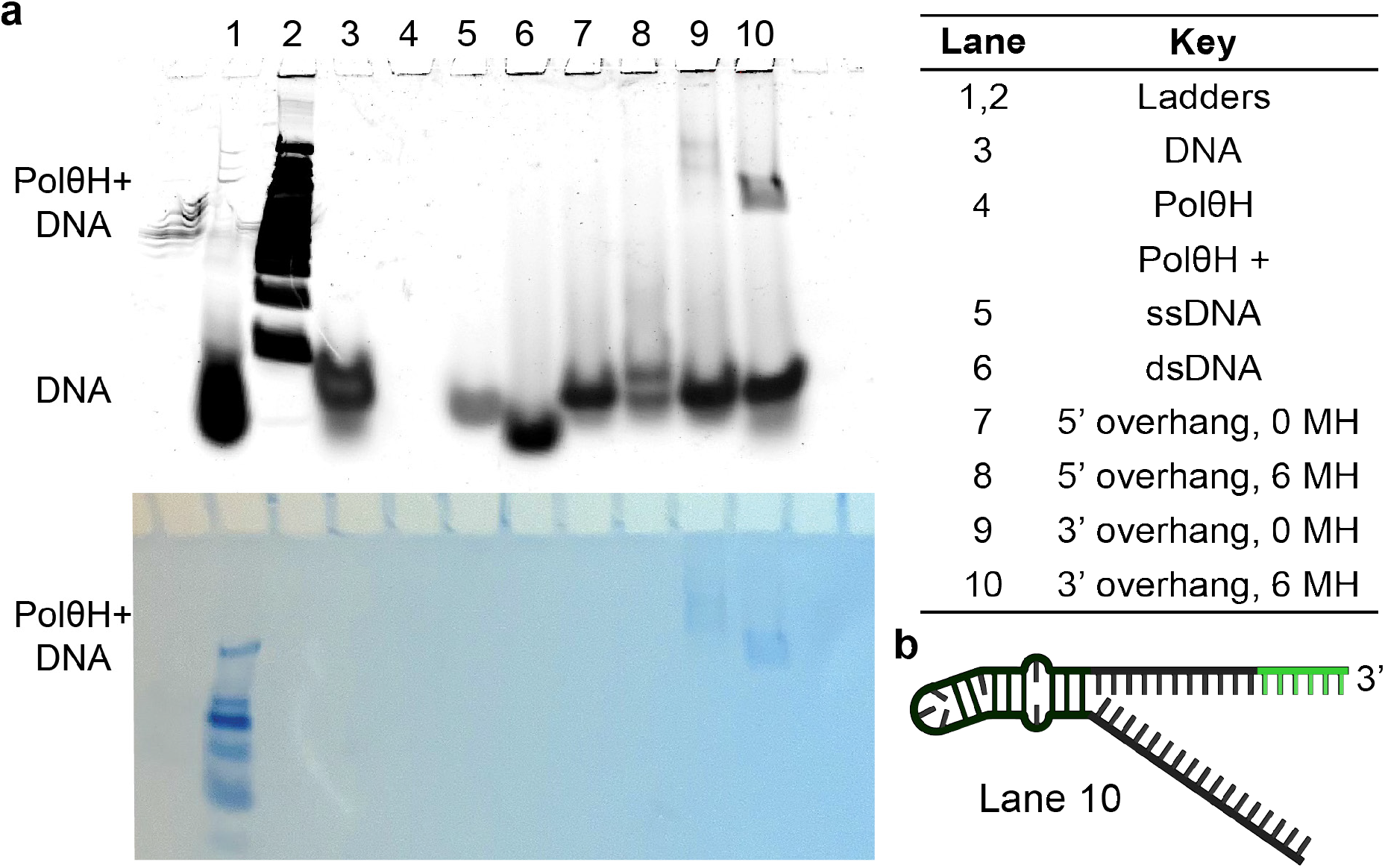
Native PAGE screening for suitable substrates for PolθH-DNA cryo-EM studies. (**a**) An example gel from Native PAGE screening for DNA substrates that bind PolθH, consecutively stained with Diamond DNA stain (top) and Coomassie protein stain (bottom). Gel migration of each species is labeled on the left, and the key is on the right. MH = microhomology. (**b**) The stem-loop DNA species pursued for structural studies, with 6 bp of self-complementary microhomology (green) at the end of the 3’ overhang.

**Supplementary Figure 3.**
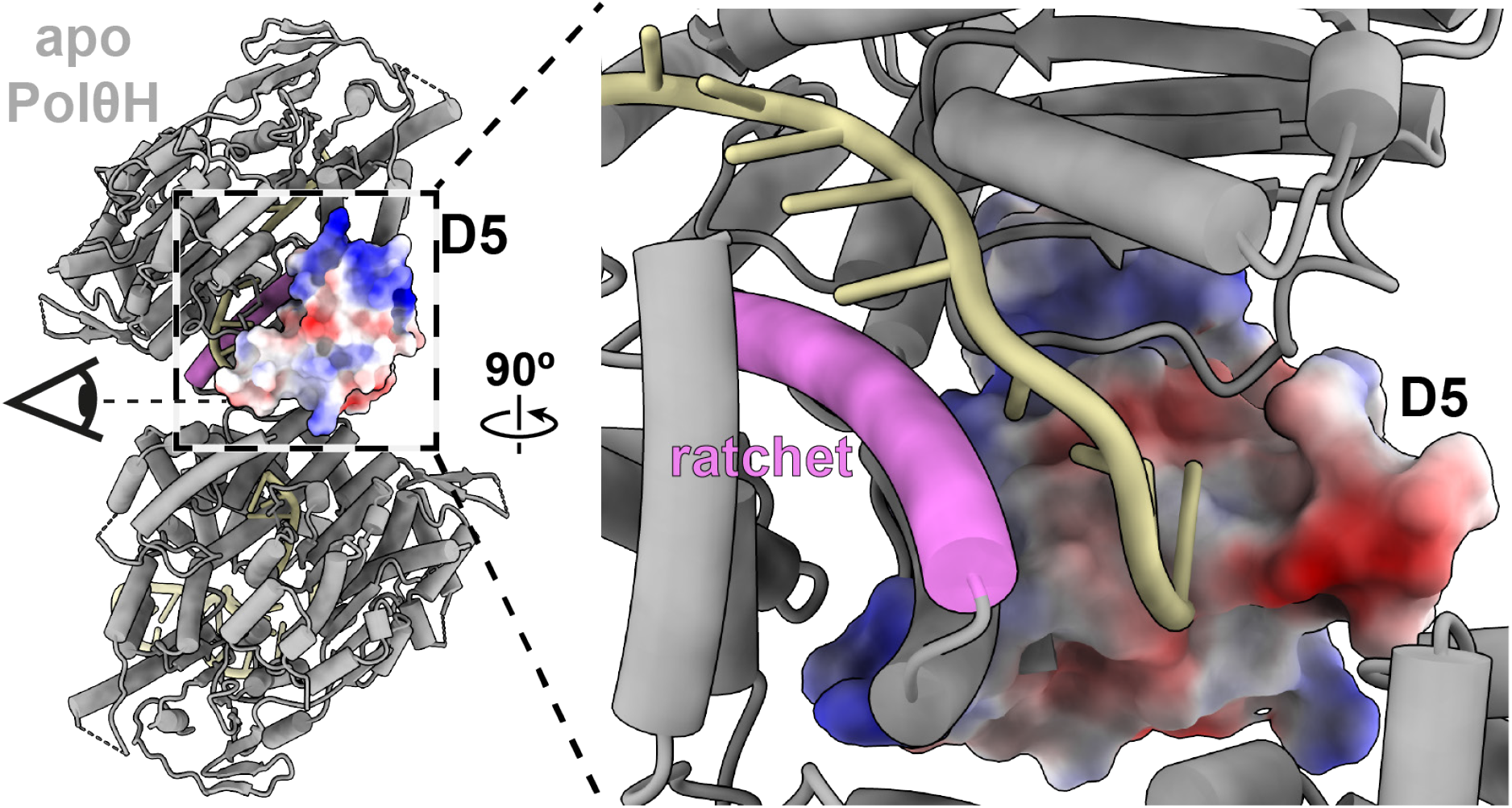
3’ ssDNA encounters a negatively charged patch on the C-terminal helix of D5 at the PolθH DNA tunnel exit. DNA from the PolθH-DNA searching model is merged with the apo PolθH dimer model. The D5 surface of one PolθH protomer is colored by electrostatic potential, and the ratchet helix is colored pink.

**Supplementary Figure 4.**
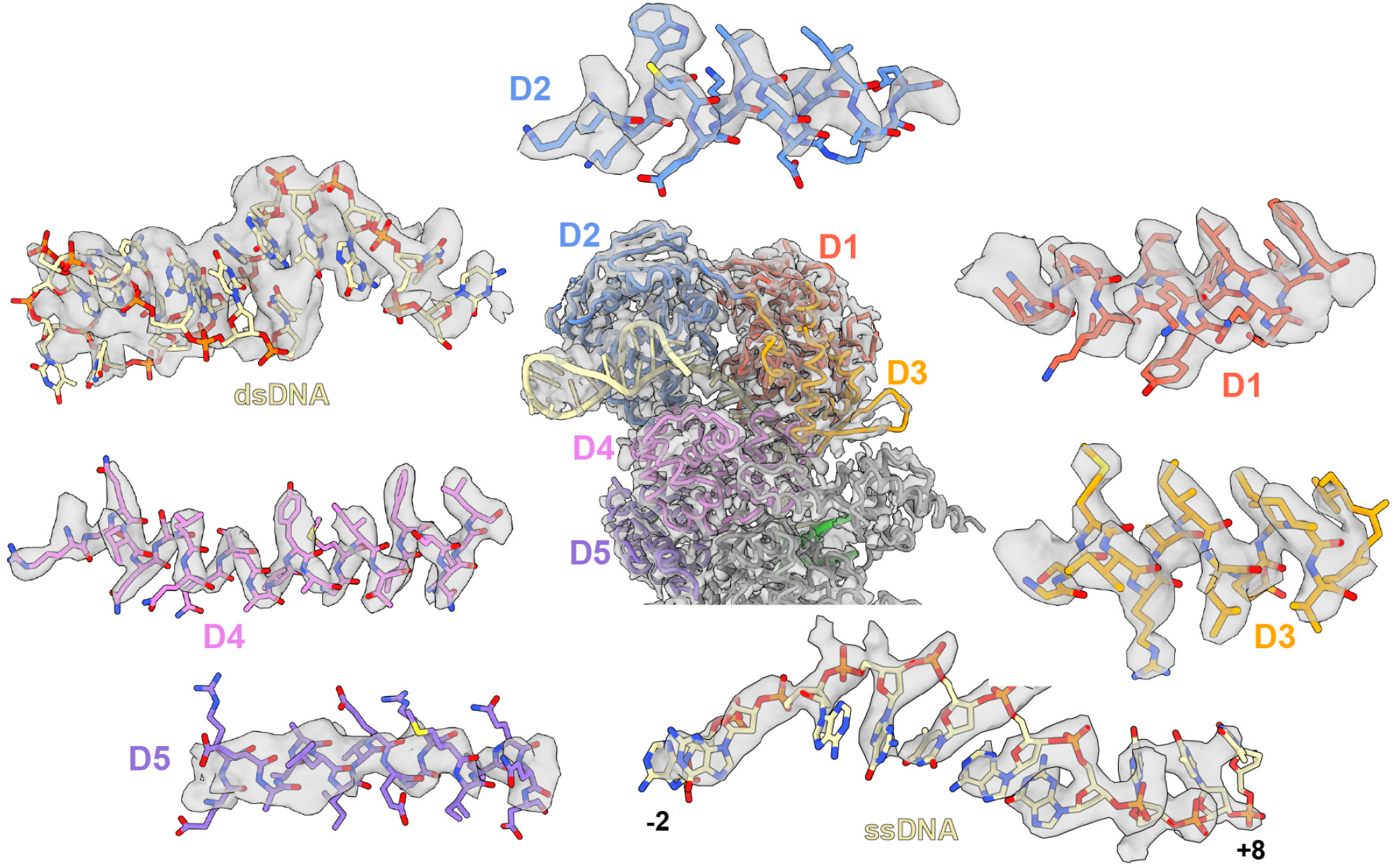
Representative cryo-EM density of PolθH. Cryo-EM map (DNA microhomology annealed state) and model of one DNA-bound PolθH protomer with domains colored, surrounded by representative map density from alpha helices in each PolθH domain. Domain 1: residues 147-161. Domain 2: residues 347-360. Domain 3: residues 529-540. Domain 4: residues 745-769. Domain 5: residues 872-890. ssDNA: bases -2 to +8. dsDNA is in the same pose as in the central protomer.

**Supplementary Figure 5.**
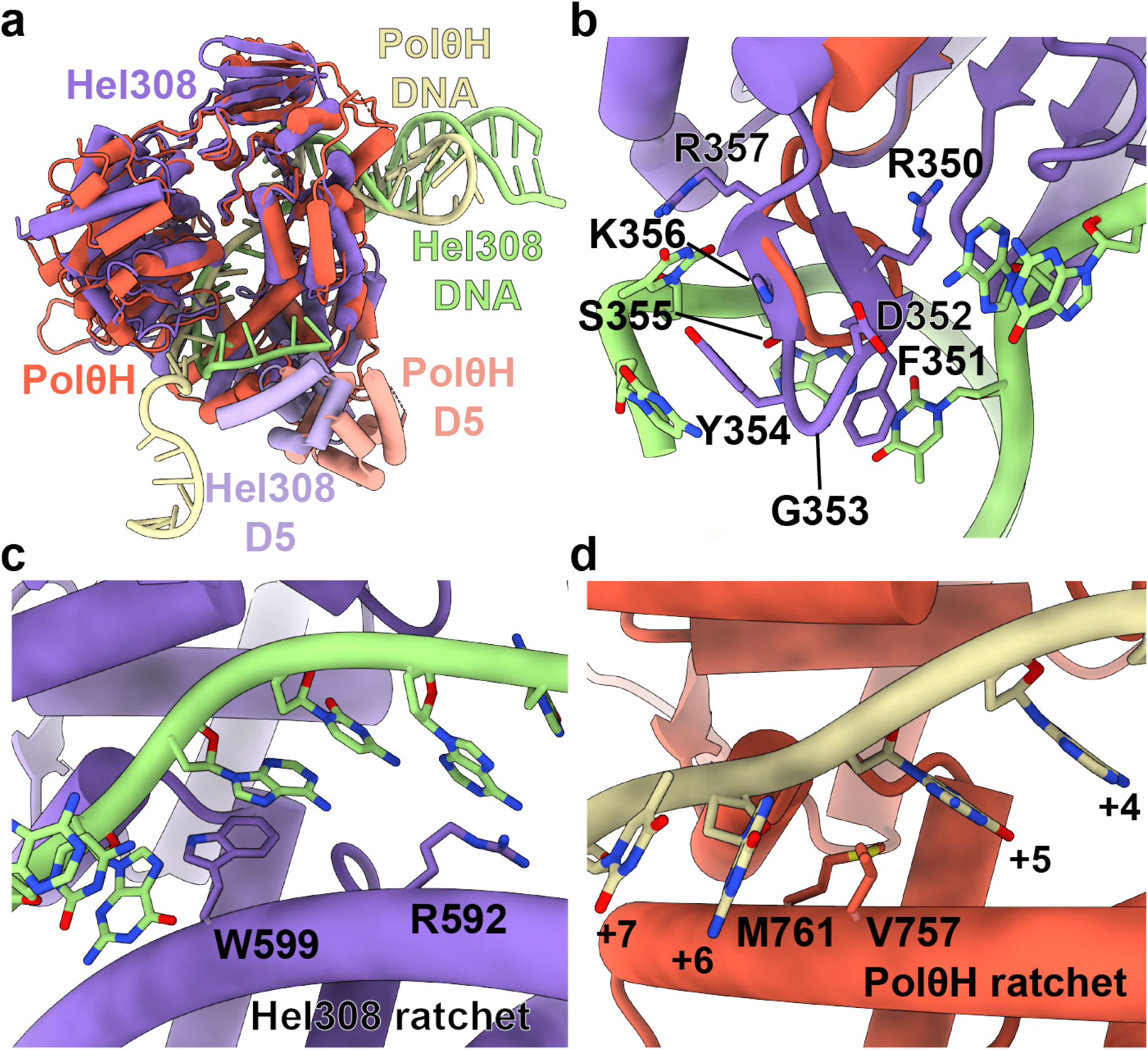
Comparison of DNA-bound structures of PolθH and *A. fulgidus* Hel308. (**a**) PolθH-DNA structure (red and yellow) overlaid with *A. fulgidus* Hel308-DNA structure (PDB: 2P6R, purple and green). (**b**) The unwinding loop of *A. fulgidus* Hel308 (purple, with residues shown) is larger than the equivalent PolθH loop (red). In the *A. fulgidus* Hel308 unwinding loop, F351 and Y354 are positioned to form pi-stacking interactions with DNA bases. Some bases in the DNA duplex have been hidden for clarity. (**c**) R592 and W599 in the *A. fulgidus* Hel308 ratchet helix are positioned to stack with DNA. (**d**) V757 and M761 in the PolθH ratchet helix wedge between DNA bases +5 and +6.

**Supplementary Figure 6.**
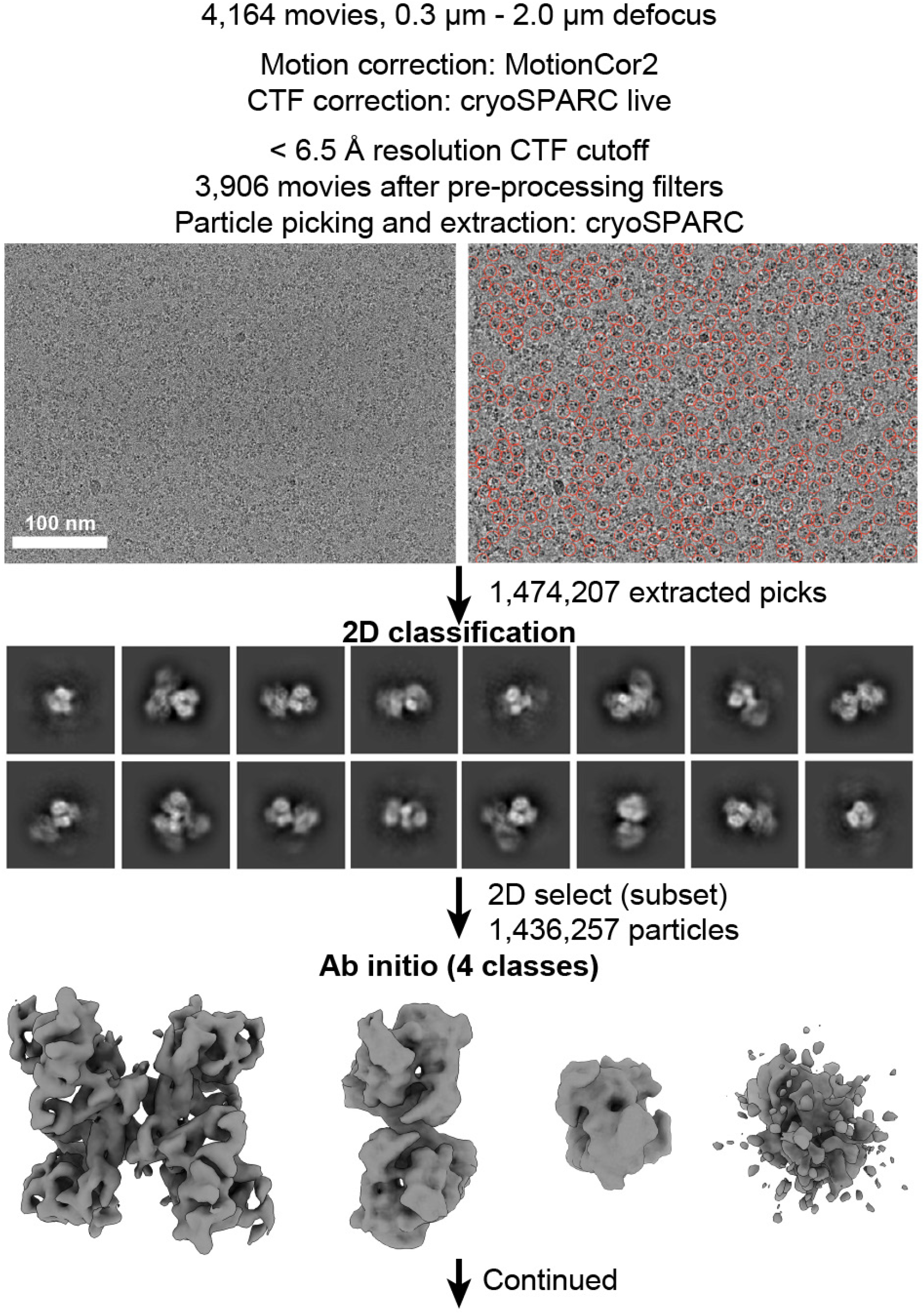
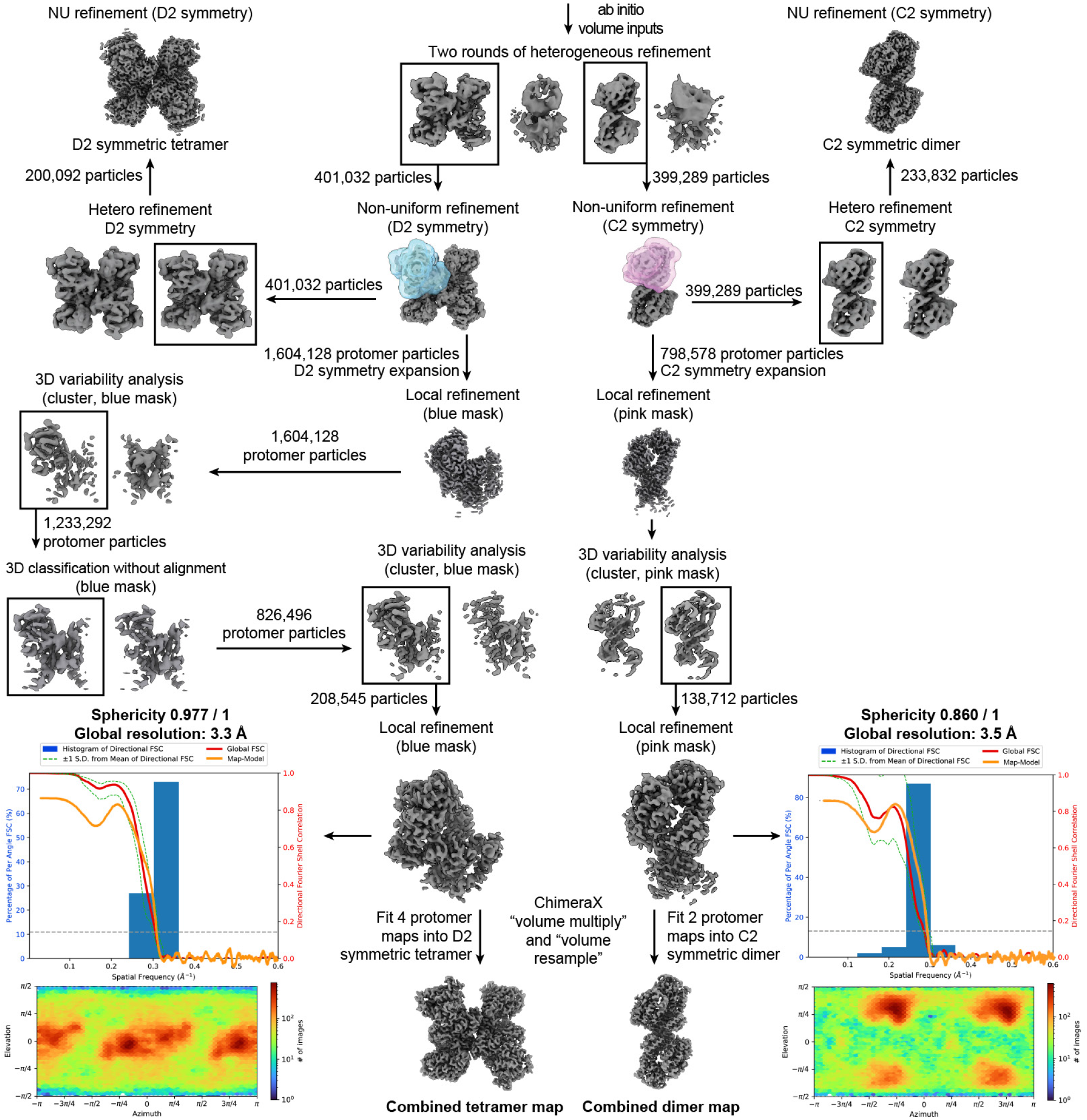
Apo PolθH cryo-EM image analysis workflow. Representative processing workflow described for apo PolθH. CTF correction was performed in cryoSPARC live, and micrographs were exported to cryoSPARC for particle picking, extraction, and further processing based on particle presence in 2D and map homogeneity in 3D as described in the methods. For 2D and 3D sorting jobs, representative classes are shown. In this scheme, initial 3D models were created Ab initio from particles and then utilized as volume inputs for iterative heterogeneous refinement, which separated tetrameric and dimeric PolθH particles. Non-uniform refinement was performed with symmetry imposed for both species to produce symmetric reconstructions. In addition, one protomer from each species was masked and symmetry expanded, subject to 3D variability analysis and/or 3D classification without alignment, and a consensus protomer of both species was locally refined. This yielded final protomer reconstructions both shown adjacent to an angular distribution plot and a 3DFSC histogram with the map-to-model FSC overlaid. For each final “combined” map, the locally refined protomer was multiplied and fit to the appropriate symmetric map.

**Supplementary Figure 7.**
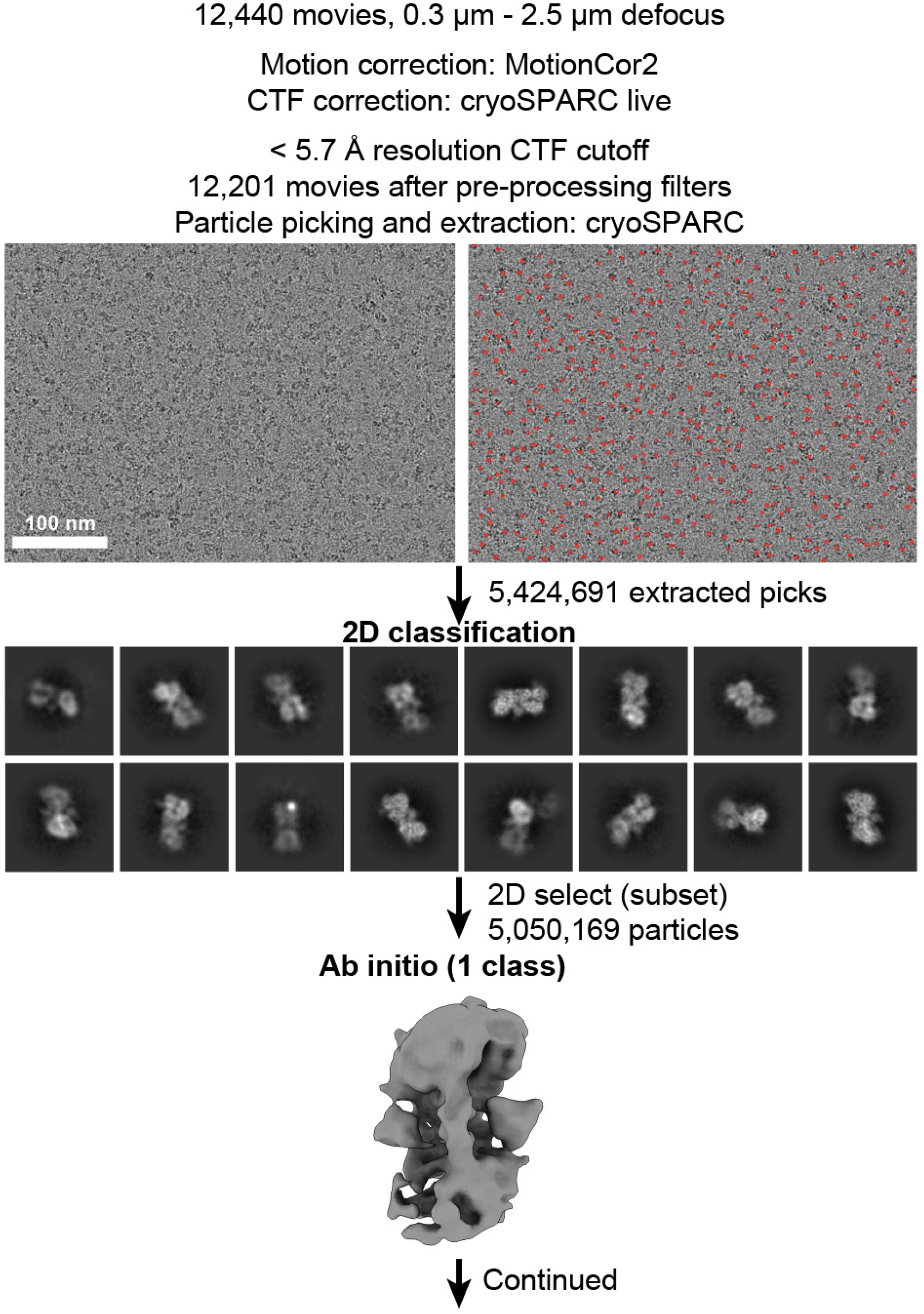
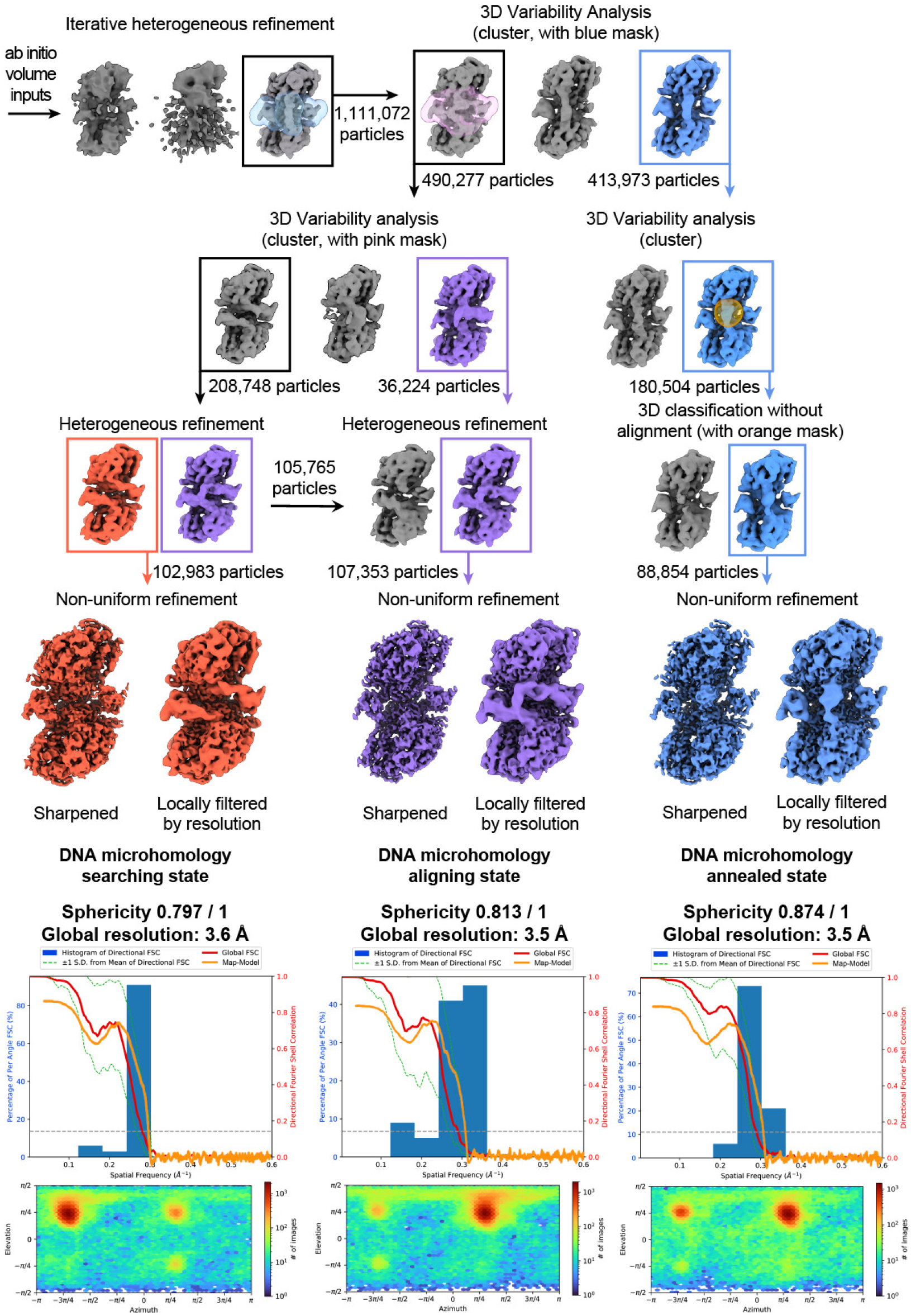
PolθH-DNA cryo-EM image analysis workflow. Representative processing workflow described for DNA-bound PolθH. CTF correction, particle picking, and particle extraction were performed in cryoSPARC live. Particles were exported to cryoSPARC for further processing based on particle presence in 2D and map homogeneity in 3D as described in the methods. For 2D and 3D sorting jobs, representative classes are shown. In this scheme, an initial 3D model was created Ab initio from particles and then utilized as a volume input for iterative heterogeneous refinement. Particles with apparent DNA density were subject to iterative 3D variability analysis with a mask encompassing the most heterogeneous regions to eliminate particles with no domain 5 density and poor DNA density. Particles were sorted into three groups based on DNA conformation (colored) and subject to non-uniform refinement to produce final reconstructions of PolθH in three different DNA pairing states, each above an angular distribution plot and a 3DFSC histogram with the map-to-model FSC overlaid. As some DNA density disappears upon map sharpening, a locally filtered map is also shown for each reconstruction.

**Supplementary Figure 8.**
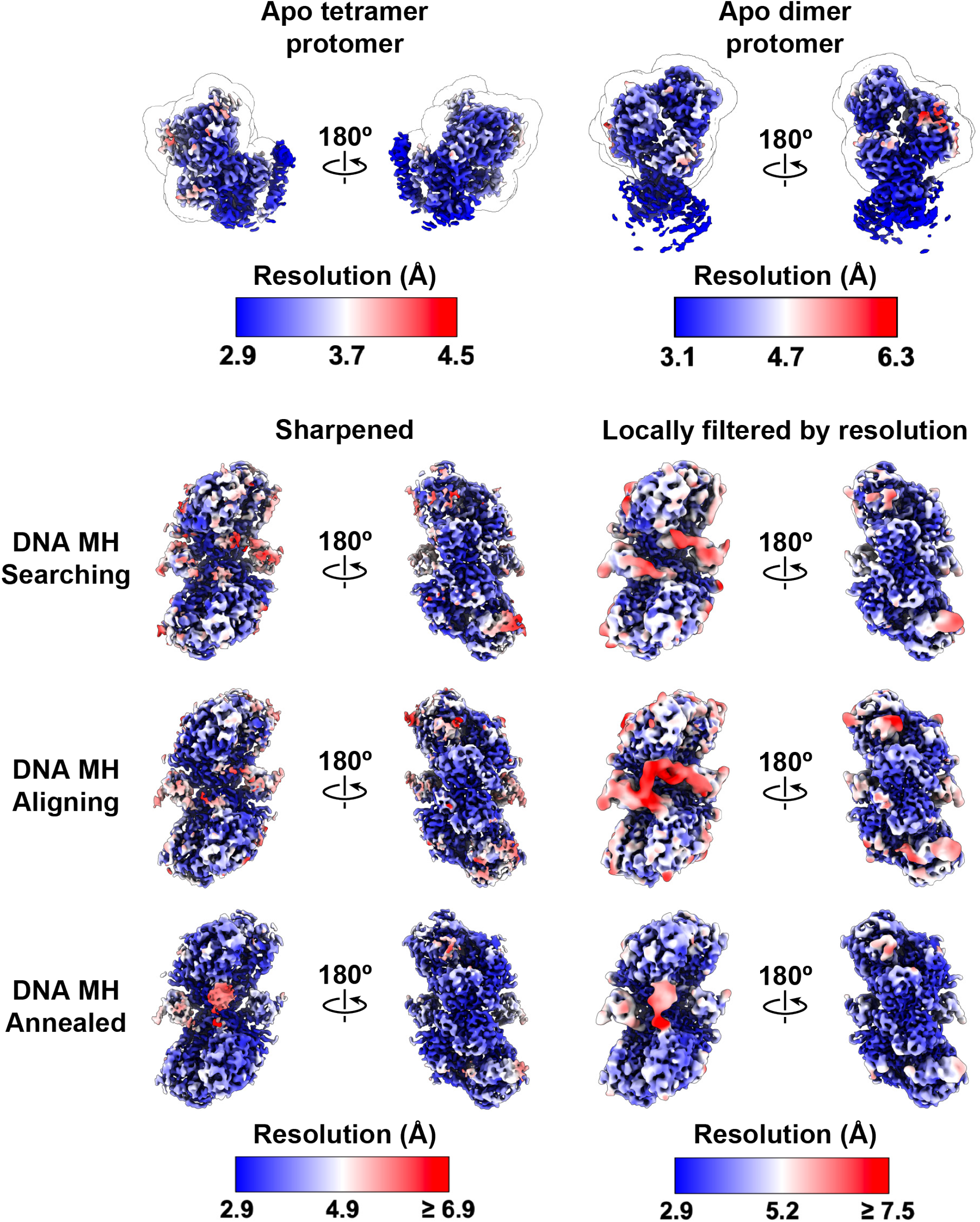
Local resolution representations of published reconstructions. Gallery of cryo-EM maps reported in this manuscript are shown colored according to their local resolution estimations. One locally refined protomer is shown for each apo map, with the mask used for local refinement outlined around it.

**Supplementary Figure 9.**
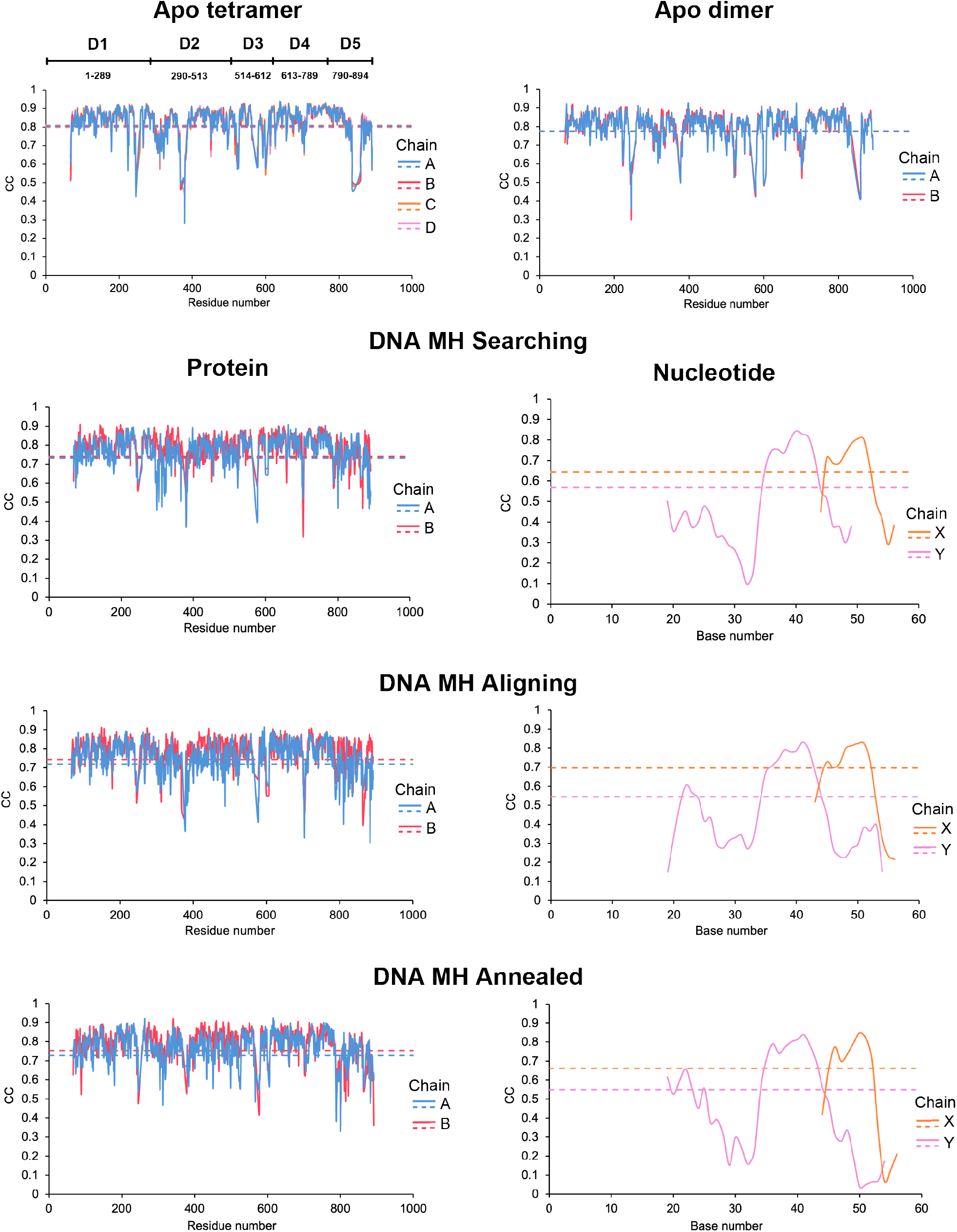
Per-residue CC plots for published models. Fitted models were analyzed with PHENIX and their per-residue correlation coefficients (CC) are plotted as compared with their respective EM reconstructions. Solid lines represent per-residue CC, and dashed lines represent average CC for the chain. At the top, PolθH domain boundaries are described relative to the plots.

**Supplementary Table 1.**
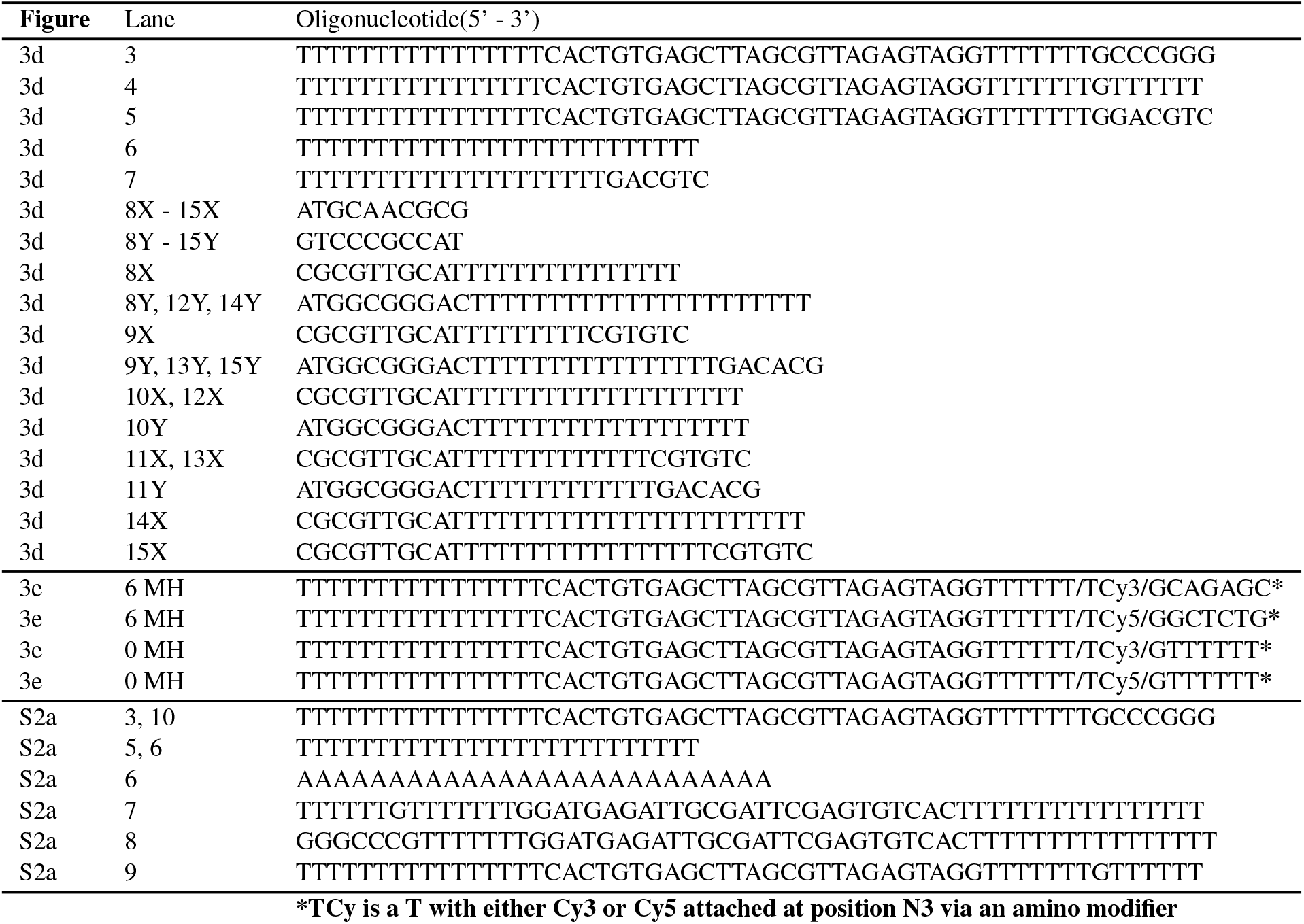
DNA oligonucleotides used in this study.

**Supplementary Table 2.** PolθH Apo and DNA-bound cryo-EM data collection, analysis, and modeling statistics.

|  | Apo Tetramer | Apo Dimer | DNA Searching | DNA Aligning | DNA Annealed |
| --- | --- | --- | --- | --- | --- |
| PDB ID | 9BH6 | 9BH7 | 9BH8 | 9BH9 | 9BHA |
| EMDB ID | EMD-44534 | EMD-44535 | EMD-44536 | EMD-44537 | EMD-44538 |
| Data Collection |  |  |  |  |  |
| Microscope | Titan Krios |  | Titan Krios |  |  |
| Camera | K3 BioQuantum |  | K3 BioQuantum |  |  |
| Camera mode | Counting |  | Counting |  |  |
| Magnification (nominal/at detector) | 105,000/60,024 |  | 105,000/60,024 |  |  |
| Voltage (kV) | 300 |  | 300 |  |  |
| Data acquisition software | Leginon |  | Leginon |  |  |
| Exposure navigation | image shift to 54 |  | image shift to 63 |  |  |
| Total electron exposure (e<sup>-</sup>/Å<sup>2</sup>) | 50 |  | 50 |  |  |
| Exposure rate (e<sup>-</sup>/pixel/sec) | 38.8 |  | 24.8 |  |  |
| Frame length (ms) | 30 |  | 30 |  |  |
| Number of frames | 30 |  | 47 |  |  |
| Pixel size (Å/pixel)* | 0.833 |  | 0.833 |  |  |
| Defocus range (μm) | -0.3 to -2.0 |  | -0.3 to -2.5 |  |  |
| Slit Width (eV) | N/A |  | 20 |  |  |
| Micrographs collected (no.) | 3,906 |  | 12,440 |  |  |
| Data analysis |  |  |  |  |  |
| Total extracted picks (no.) | 1,474,207 |  | 5,424,691 |  |  |
| Particles used for 3D (no.) | 1,436,257 |  | 5,050,169 |  |  |
| Final Particles (no.) | 208,545* | 130,712* | 102,983 | 107,353 | 88,854 |
| Symmetry | D2 | C2 | C1 | C1 | C1 |
| Resolution (global) (Å) |  |  |  |  |  |
| FSC 0.5 (unmasked / masked) | 4.4* / 3.7* | 4.7* / 4.0* | 5.0 / 4.0 | 4.6 / 4.0 | 4.5 / 3.7 |
| FSC 0.143 (unmasked / masked) | 3.8* / 3.3* | 4.1* / 3.5* | 4.0 / 3.5 | 4.0 / 3.4 | 3.8 / 3.3 |
| Local resolution range (Å) | 2.9* - 4.4* | 3.1* - 6.3* | 3.1 - 7.4 | 3.0 - 6.8 | 2.9 - 7.7 |
| 3DFSC Sphericity (%) | 97.7* | 86.0* | 79.7 | 81.3 | 87.4 |
| Map sharpening *B*-factor (Å<sup>2</sup>) | -90* | -112* | -102.1 | -102.6 | -108.7 |
| Model composition |  |  |  |  |  |
| Chains | 4 | 2 | 4 | 4 | 4 |
| Non-hydrogen atoms | 24,388 | 12,166 | 13,051 | 13,155 | 13,270 |
| Protein residues | 3,096 | 1,544 | 1,574 | 1,574 | 1,586 |
| Nucleotides | 0 | 0 | 44 | 50 | 49 |
| Model Refinement* |  |  |  |  |  |
| Refinement package | ISOLDE/Phenix | ISOLDE/Phenix | ISOLDE/Phenix | ISOLDE/Phenix | ISOLDE/Phenix |
| MapCC (volume/mask) | 0.82 / 0.81 | 0.77 / 0.78 | 0.74 / 0.76 | 0.74 / 0.77 | 0.74 / 0.77 |
| B-factors (Å<sup>2</sup>) |  |  |  |  |  |
| Protein residues | 54.43 | 55.33 | 69.39 | 67.19 | 53.26 |
| Nucleotides | N/A | N/A | 33.75 | 42.19 | 35.80 |
| R.m.s. deviations |  |  |  |  |  |
| Bond lengths (Å) | 0.003 | 0.003 | 0.003 | 0.003 | 0.003 |
| Bond angles (°) | 0.614 | 0.634 | 0.640 | 0.680 | 0.654 |
| Validation |  |  |  |  |  |
| Map-to-model, FSC 0.5 | 3.5* | 3.7* | 3.8 | 3.7 | 3.7 |
| Ramachandran plot |  |  |  |  |  |
| Favored (%) | 99.38 | 99.21 | 99.29 | 99.35 | 99.42 |
| Allowed (%) | 0.49 | 0.66 | 0.71 | 0.65 | 0.51 |
| Outliers (%) | 0.13 | 0.13 | 0.00 | 0.00 | 0.06 |
| MolProbity score | 1.44 | 1.58 | 0.90 | 0.93 | 0.92 |
| Clashscore | 1.44 | 1.58 | 1.53 | 1.75 | 1.70 |
| Poor rotamers (%) | 0.00 | 0.00 | 0.15 | 0.07 | 0.00 |
| CaBLAM outliers (%) | 0.30 | 0.40 | 0.52 | 0.39 | 0.45 |
| EMRinger score<sup>64</sup> | 3.48 | 2.24 | 2.33 | 3.06 | 2.78 |
| Qscore | 0.53 | 0.48 | 0.46 | 0.48 | 0.47 |

\* refers to asymmetric units after symmetry expansion (single protomer)

